# Hypothalamic recurrent inhibition regulates functional states of stress effector neurons

**DOI:** 10.1101/2025.09.03.673779

**Authors:** Aoi Ichiyama, Samuel Mestern, Tamas Fuzesi, Brian L. Allman, Wilten Nicola, Jaideep Bains, Lyle Muller, Wataru Inoue

## Abstract

Stress triggers rapid and reversible shifts in vital physiological functions from homeostatic operation to emergency response. However, the neural mechanisms regulating such functional stress states remain poorly understood. Here we identify a novel recurrent inhibitory circuit governing functional states of key stress regulatory neurons: corticotropin-releasing hormone (CRH) neurons in the hypothalamic paraventricular nucleus (PVN). Microendoscopic calcium imaging in freely behaving mice revealed synchronized low-activity state at baselines and a reversible high-activity state during mild stress. Ensemble analysis indicated increased dimensionality of network dynamics during high-activity state. Computational modeling of calcium ensemble data, together with independent modeling of single-unit CRH_PVN_ neurons spiking dynamics, converged to show that recurrent inhibition is a key circuit motif for stress-induced functional state transitions. Guided by model predictions, chemogenetic manipulations of PVN-projecting GABAergic neurons (^PVN→^GABA) revealed their roles in constraining CRH_PVN_ neurons to low-activity state at baselines via a prolonged feedback inhibition. Unexpectedly, slow CRHergic excitation was dispensable for driving this prolonged feedback, whereas glutamatergic transmission predominated at CRH→^PVN→^GABA excitatory synapses.

Incorporating these findings, we refined our computational model to include fast excitation and slow inhibition, yielding new predictions for circuit operation. Together, our results establish recurrent inhibition as a fundamental circuit motif controlling CRH_PVN_ neurons functional states and highlight the value of iterative experiment–model integration in advancing understanding of neural circuits functions.

## INTRODUCTION

A threat to well-being triggers a range of physiological and behavioural changes, including hormone release, elevated heart rate, escape and fight, collectively termed as the stress response^1^. However, how the brain generates, maintains and resolves functional stress states that underlie these pleiotropic changes remain largely unknown.

Many aspects of stress are governed by the hypothalamus, an evolutionarily-conserved brain region crucial for homeostasis regulation and survival behaviours. Among hypothalamic neurons, corticotropin-releasing hormone (CRH) neurons in the paraventricular nucleus of the hypothalamus (PVN) are classically known as central regulator of the hormonal stress response. However, emerging evidence uncovers a broader role for CRH_PVN_ neurons in multifaceted stress response. For example, optogenetic or chemogenetic stimulation of CRH_PVN_ neurons induces a repertoire of defensive behaviors^2–4^, provokes negative emotions^5,6^, promotes wakefulness^7^, and social stress transmission^8^. Conversely, inhibiting these neurons reverse these changes during stress, causally linking CRH_PVN_ neurons activity with pleiotropic stress responses. A key open question is what are neural circuits that govern the dynamic functional states of CRH_PVN_ neurons.

*In vivo*, individual neurons constantly receive bombardments of excitatory and inhibitory synaptic inputs from the network they are embedded in^9,10^, and such network dynamics link functional states of circuits and constituent single neurons^11^. Using single-unit recording from anesthetized mice, we recently revealed that, in response to stress stimuli, individual CRH_PVN_ neurons profoundly change their firing patterns from rhythmic low-activity to tonic high-activity states^12^. Subsequent computational modeling found that these *in vivo* firing dynamics can be effectively recapitulated in recurrent inhibitory circuits where a loss of recurrent inhibition mediates the transitions from the low-to high-activity states. Importantly, creating a neural model enables us to abstract complex *in vivo* phenomenon into a concise representation—for example recurrent inhibition circuit motif—and demonstrate that its dynamic activities can support functional states transitions of CRH_PVN_ neurons. Such a data-driven model, in turn, serves as working hypotheses, providing explicit predictions that can then be experimentally interrogated by direct manipulations to establish causality, or reveal key biological features for iterative refinements of the model.

Here, guided by the circuit model, we experimentally examined recurrent inhibitory circuits using miniature microscope calcium imaging in freely moving mice, single-unit recordings, and chmogenetic, optogenetic and pharmacological circuit manipulations. We show ubiquitous evidence for the recurrent inhibitory circuit in mediating functional state transitions of CRH_PVN_ neurons by stress and offer new insights into the mechanisms that regulate functional stress states.

## RESULTS

### Stress elicits a transition from low-to high-activity states of CRH_PVN_ neurons

We used *in vivo* miniaturized microscopy to characterize CRH_PVN_ neurons activity dynamics in freely behaving mice. A GRIN lens was implanted above GCaMP6f-expressing CRH_PVN_ neurons to monitor intracellular Ca²⁺ dynamics (Fig. 1A-D). In the homecage, CRH_PVN_ neurons exhibited sporadic activity presenting their baseline activity. The transfer to a novel environment (a clean plastic chamber without bedding) dramatically increased the overall activity (Fig. 1E-G). Specifically, in the homecage, CRH_PVN_ neurons exhibited sporadic activity that increased almost seven-fold during exposure to a novel context (Fig. 1H). Furthermore, the vast majority of the CRH_PVN_ neurons responded with a significant increase in detected events (Fig. 1I). The summed activity of individual neurons in the novel environment revealed a population-level tonic increase, which persisted throughout the exposure (Fig. 1G). This high-activity state was absent upon returning to the homecage, where activity returned to the pre-transfer low-activity states (Fig. 1E-I), indicating that the activity state is fully reversible and is reflecting animals’ response to a mildly stressful, novel environment.

**Fig. 1.**
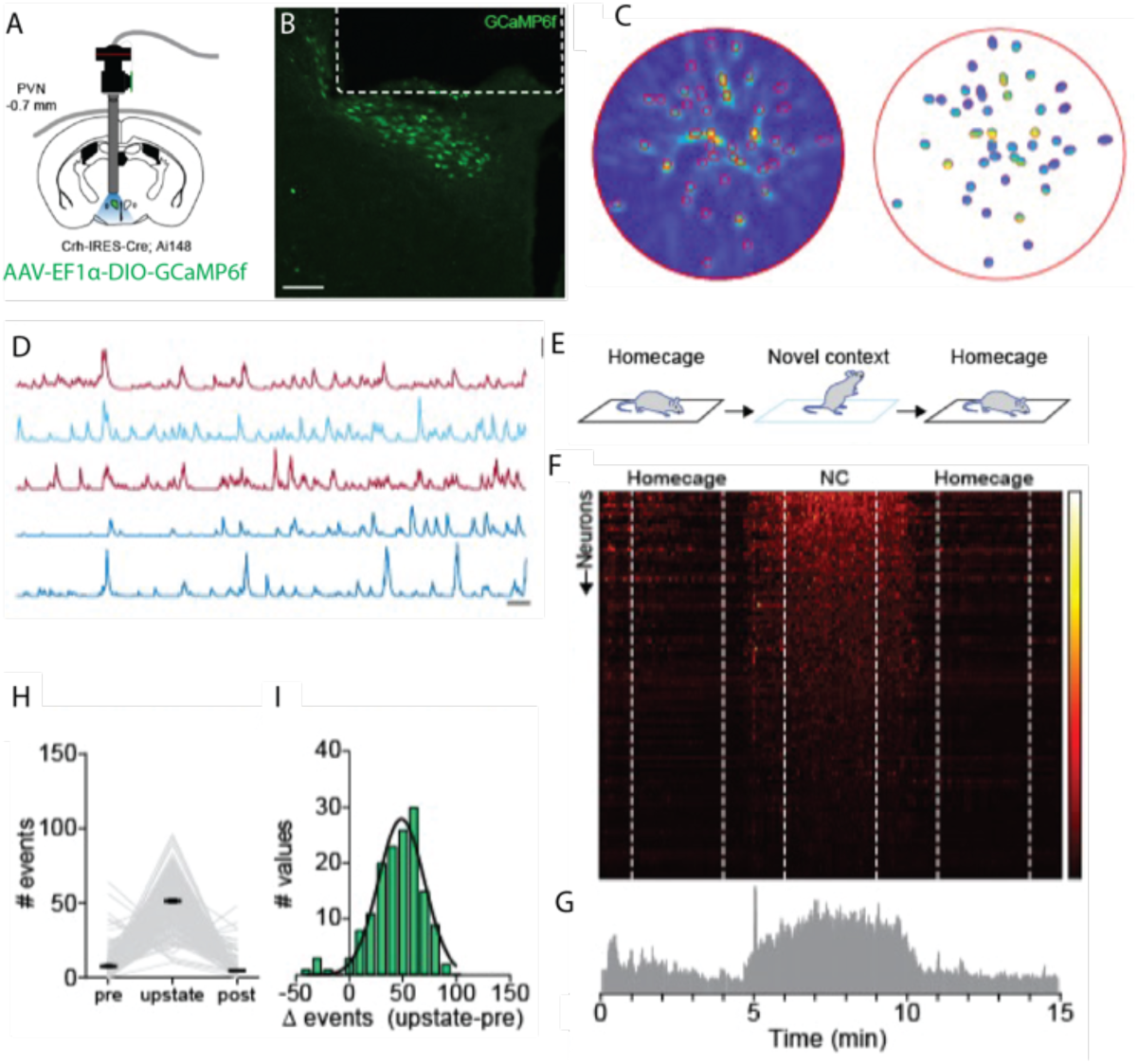
Novel context elicits a transition from low-to high-activity states of CRH^PVN^ neurons. **A,** Schematic map shows the GRIN lens implantation above the PVN of CRH-IRES-Cre mice injected with Cre dependent AAV-DIO-GCaMP6s. **B,** Confocal image shows expression of GCaMP6s (green) in the PVN and GRIN lens position (dotted line). **C,** Miniature microscope field of view and identified units using the Min1pipe analysis. **D,** Sample traces of individually identified units. **E,** Schematic experimental design of exposure to a novel context (NC). **F,** Normalized heat map shows the activity of every identified neurons during the 15 min recording. Colour bar for intensity on the right. **G,** Summarized activity in 1-sec bins shows the distribution of PVN CRH activity. **H,** Number of events detected from the identified units when the animal is in its homecage (pre), placed into the novel context (high-activity state) and returned to the homecage afterwards (post) (pre: 7.62 ± 0.83 events; upstate: 51.18 ± 1.41 events; post: 4.43 ± 0.63 events; pre vs upstate, p<0.0001; upstate vs post, p<0.0001; pre vs post, p=0.1402; n=152, repeated measures 1-way ANOVA F(2,302)=536.5, P<0.0001, Bonferroni’s multiple comparisons test). **I,** Histogram shows the change in the number of events generated by individual neurons between the upstate and pre period. Scale bars, **b**, 20 µm; **D**, 10 s; **G,** 20 %. Data are mean ± s.e.m.

### Circuit dynamics regulate CRH_PVN_ neurons activity state transitions

To further investigate the neural ensemble dynamics underlying these reversible activity states, we applied Singular Value Decomposition (SVD) to 3-minute intervals during the baseline homecage, novel context, and post-homecage sessions (Fig. 2A-C). The first SVD feature revealed an overall increase in activity during the novel context. Notably, in the homecage, synchronized activity was observed, represented by Ca^2+^ spikes in the second dominant SVD feature across all animals (Fig. 2C, Extended Data Fig. 1). By contrast, during the novel environment exposure, despite the overall increase in event rate, there were no synchronized population activities (Fig. 2C), and the second SVD feature displayed a continuous distribution of positive and negative values (Fig. 2C), consistent with the data that although some neurons show an increase in their Ca^2+^ signal during the home-cage bursts, others show a decrease (Fig. 1I, Extended Data Fig. 1). Interestingly, the synchronized population activities during the pre- and post-homecage sessions did not appear to result from local interactions between nearby neurons (Fig. 2D, Extended Data Fig. 1). A comparison of Pearson correlation coefficients between nearest neighbors and a neuron-shuffled distribution showed no statistically significant difference (Fig. 2E, Extended Data Fig. 1). Additionally, reconstructing the data using increasingly larger numbers of SVD features revealed that more features were required to reach a fixed level of reconstruction error (10%) in the novel context than in either the pre- or post-homecage sessions (Fig. 2F). This suggests that the high-activity state represents not only increased overall activity but also increased dimensionality in the underlying neural ensemble dynamics.

**Fig. 2.**
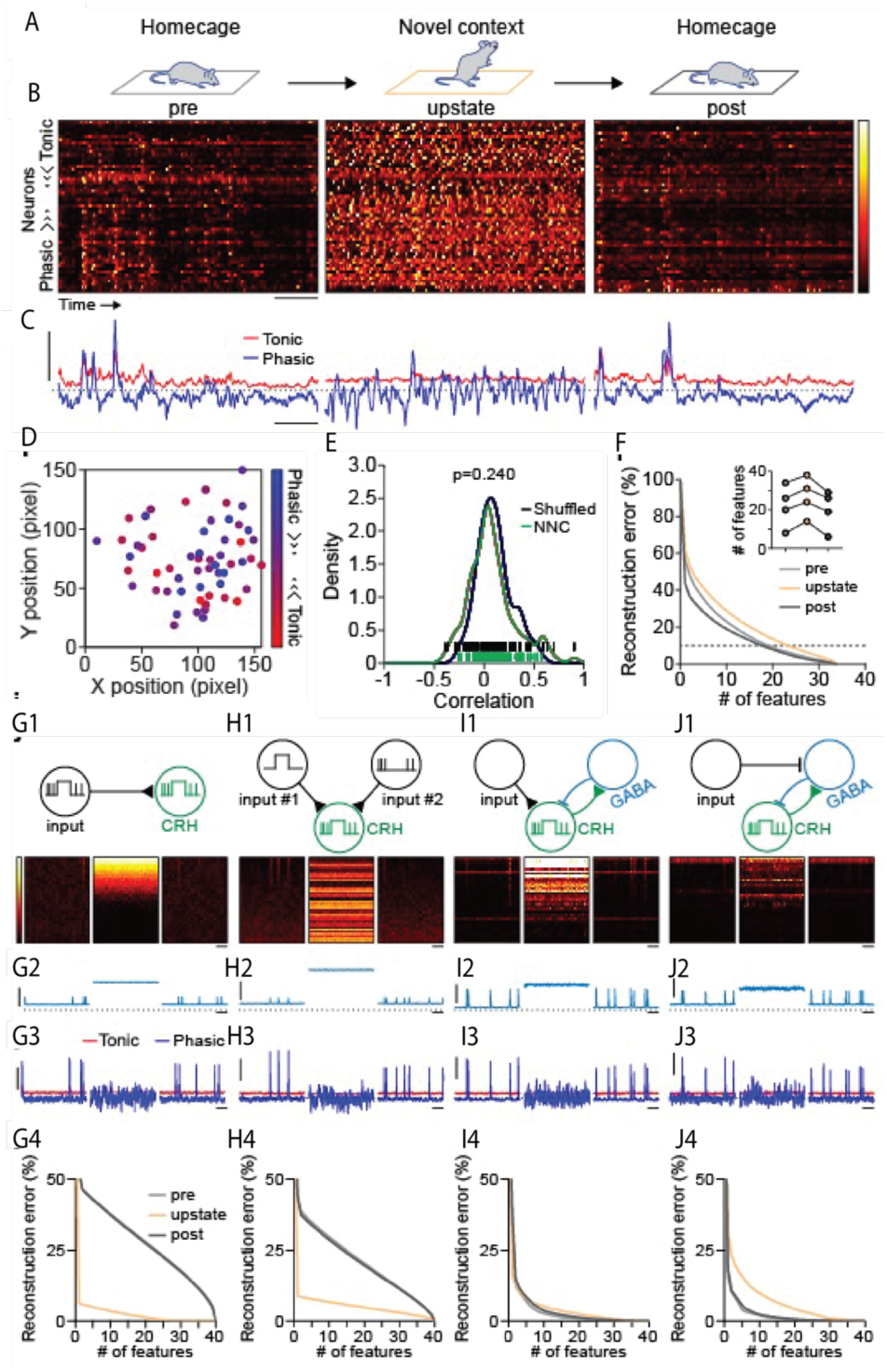
Local network dynamics contribute to the increased desynchronized activity of CRHPVN neurons during the high-activity state. **A**, Schematic experimental design of exposure to NC. **B**, 3 minute episodes of miniature microscopy recordings when the animal is in its homecage (pre), placed into the novel context (high-activity state) and returned to the homecage afterwards (post). The neurons are sorted based on the coefficients projected onto the second basis function in the singular value decomposition of neural data (shown in **C**). Colour bar for intensity is on the right. **C,** The first two features in the singular value decomposition (red, tonic; blue, phasic). **D**, Topographical map of recorded neurons. The colour represents the strength of the phasic (blue) and tonic (red) components of the coefficient. **E**, The distribution of Pearson correlation coefficients between nearest neighbour ROIs for all 4 animals for data (green) and shuffled neuron indices (black). F, Graphs demonstrate the dimensionality of the calcium signal during the pre, upstate and post phase in one animal. Inset, The number of features necessary to allow only 10% reconstruction error during the three phase, roughly indicative of the dimensionality of the neural responses. **G**, A model where the CRHPVN neurons track inputs to create the high-activity state and the home-cage bursts. The schematic (**G1**), with the simulated network output (**G2**), the singular value decomposition of the synthetic calcium signal (**G3**) and the dimensional analysis (as in **I**) (**G4**). **H**, A second tracking model where the CRHPVN neurons track the inputs from two sources, one-responsible for the high-activity state and one responsible for the home cage bursts. **I**, A more detailed model consisting of inputs onto a coupled network model consisting of subpopulations of CRHPVN and GABAergic neurons. **J**, An identical model as in **I,** only the inputs inhibit the GABAergic neurons, triggering a disinhibitory response to activate the upstate and bursts in the CRHPVN neurons. Scale bars, **B**, 30 s; **C**, 0.1; **G2-J2**, 5, 30 s; **G3-J3**, 0.1, 30 s.

**Extended Data Fig. 1.**
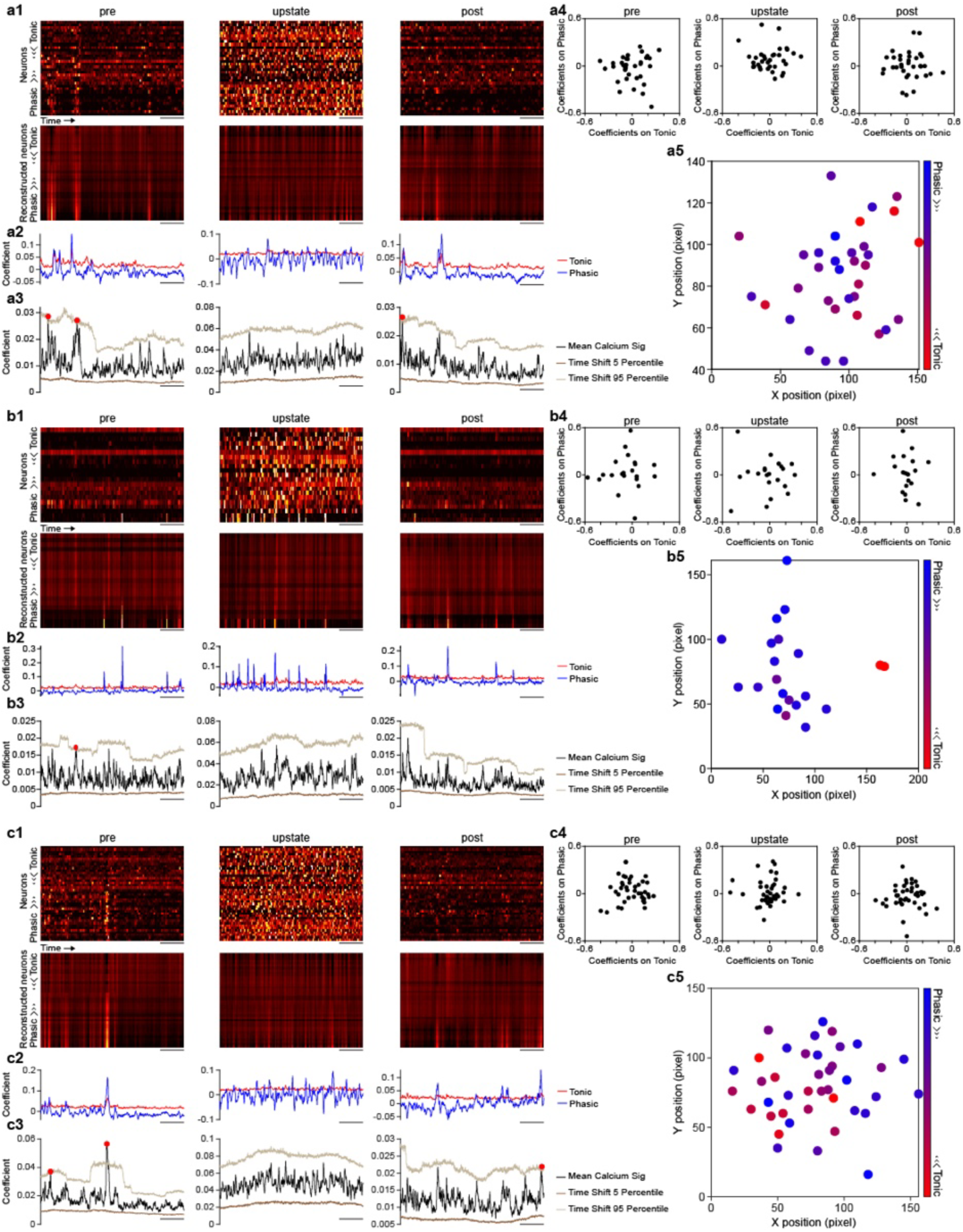
Neuronal activity analyses of additional animals. **a1**, The normalized calcium traces (see Methods) for the pre, novel context, and post homecage sessions (top). Computational reconstruction of neuronal activity using the tonic and phasic features in the pre-section (bottom). **a2**, Time aligned singular value basis functions (first two dominant values) are plotted. **a3**, Synchrony analysis demonstrating statistically significant events at a Bonferroni-corrected p=0.05 level (red circle indicates significant peaks). **a4**, Coefficients in the for the first two dominant singular value basis functions (tonic and phasic elements). **a5**, The region of interests defining a calcium trace, with the basis coefficient determining the colour (tonic to phasic). **b-c**, Identical analyses as in **a1-a5** for two additional animals. Scale bars, **a1-3**, **b1-3**, **c1-3**, 30 s.

To examine circuit motifs crucial for mediating the transitions between these low- and high-activity states, we modeled small networks of Leaky-Integrate-and-Fire (LIF) neurons of varying complexity. Initial models, which tracked a one-way incoming signal without recurrent processing (Fig. 2G-H). These are uncoupled CRH_PVN_ neuron models that receive either one input source that displays a phasic mode in the home-cage and a tonic mode in the novel context (Methods; Fig. 2G), or two separate inputs that play phasic and tonic modes independently (Fig. 2H). While both tracking models can reproduce home-cage synchronicity and the novel context high-activity state (Fig. 2G-H), the resulting dimensionality of the neural dynamics is the opposite of our experiments; the high-activity state is lower dimensional, indicating a general loss of complexity in the neural dynamics (Fig. 2G-H). To address this mismatch, we considered networks that relied on tonic input to activate higher-dimensional dynamics via recurrent connectivity, thereby increasing rather than decreasing dynamical dimension. We considered a simple coupling motif: local GABAergic neurons synapse onto CRH_PVN_ neurons, while CRH_PVN_ neurons synapse onto the GABAergic neurons (Fig. 2I). In this case, stimulating CRH_PVN_ neurons with tonic/phasic input reproduced both the novel context high-activity state and homecage low-activity state with synchronicity, but the dimensionality of the neural dynamics remained similar between the two states (Fig. 2I). However, when we suppressed GABAergic neurons, disinhibition resulted in the high-activity state and a context (Fig. 2J). These modeling results indicate that recurrent inhibition is required to account for the transitions between the low- and high-activity states of CRH_PVN_ neurons in freely moving mice.

Strikingly, recurrent inhibition was identified by our earlier single-unit electrophysiology data^12^ as a key circuit motif regulating activity state transitions of CRH_PVN_ neurons, despite major differences in the nature of neural data and experimental conditions. Specifically, miniature microscopy examined temporal correlations Ca^2+^ signals among ensembles of CRH_PVN_ neurons in freely behaving animals. On the other hand, *in vivo* electrophysiology^12^ examined temporal structure of firing patterns in single neurons recorded from anesthetized mice. Collectively, our results support ubiquitous role of recurrent inhibition in regulating stress-relevant functional state of CRH_PVN_ neurons. In the following sections, we use *in vivo* electrophysiology under anesthesia, allowing for precise control of experimental conditions, to experimentally test the causal roles of recurrent inhibitory circuit in regulating CRH_PVN_ neurons functional states.

### Spike time variability in single CRH_PVN_ neurons reflects functional states of local circuits and their stress-induced transitions

Using a high-density silicone electrode Neuropixcel^13^, we recorded CRH_PVN_ neurons that were optogenetically identified under urethane anesthesia in mice (Fig. 3A, B). As reported previously^12^, a subset of CRH_PVN_ neurons show prominent rhythmic, brief bursts intervened by long interburst silences that is characteristics of the low-activity state in baseline recording (Fig. 3C, D). This bursting phenotype, however, is heterogeneous among CRH_PVN_ neurons, and about half show little to no bursts (burst rate <0.1 Hz, Fig. 3C, D): bursting neurons tended to be located dorsal part of the PVN (Extended Data Fig. 2). Importantly, a pair of simultaneously recorded bursting and non-bursting neurons showed temporally correlated fluctuations in their firing patterns under the baseline condition (Fig. 3E, F), akin to the synchronized population activities observed in miniature microscope during low-activity states in homecage (Fig. 2A-C). To analyze these neural state dynamics, we calculated CV2, a measure of local variability of interspike intervals (ISIs). CV2 quantifies the difference between successive ISIs (i.e. Post ISI – Pre ISI; Fig 2E) relative to their mean ((Post ISI + Pre ISI)/2), providing a sensitive index of firing irregularity^14^. In essence, high CV2 values reflect greater variability in spike timing, such as during rhythmic firing, while a low CV2 value indicates more regular firing. The example pair of units shown in Fig. 3E-H exhibited highly correlated CV2 fluctuations over time (Fig. 3I). This indicates that CRH_PVN_ neurons, regardless of bursting and non-bursting firing, are influenced by shared afferent inputs and undergo coordinated state transitions. Coronally to this conclusion is that single unit firing patterns offers an effective means to infer functional states of local network involving CRH_PVN_ neurons.

**Fig. 3.**
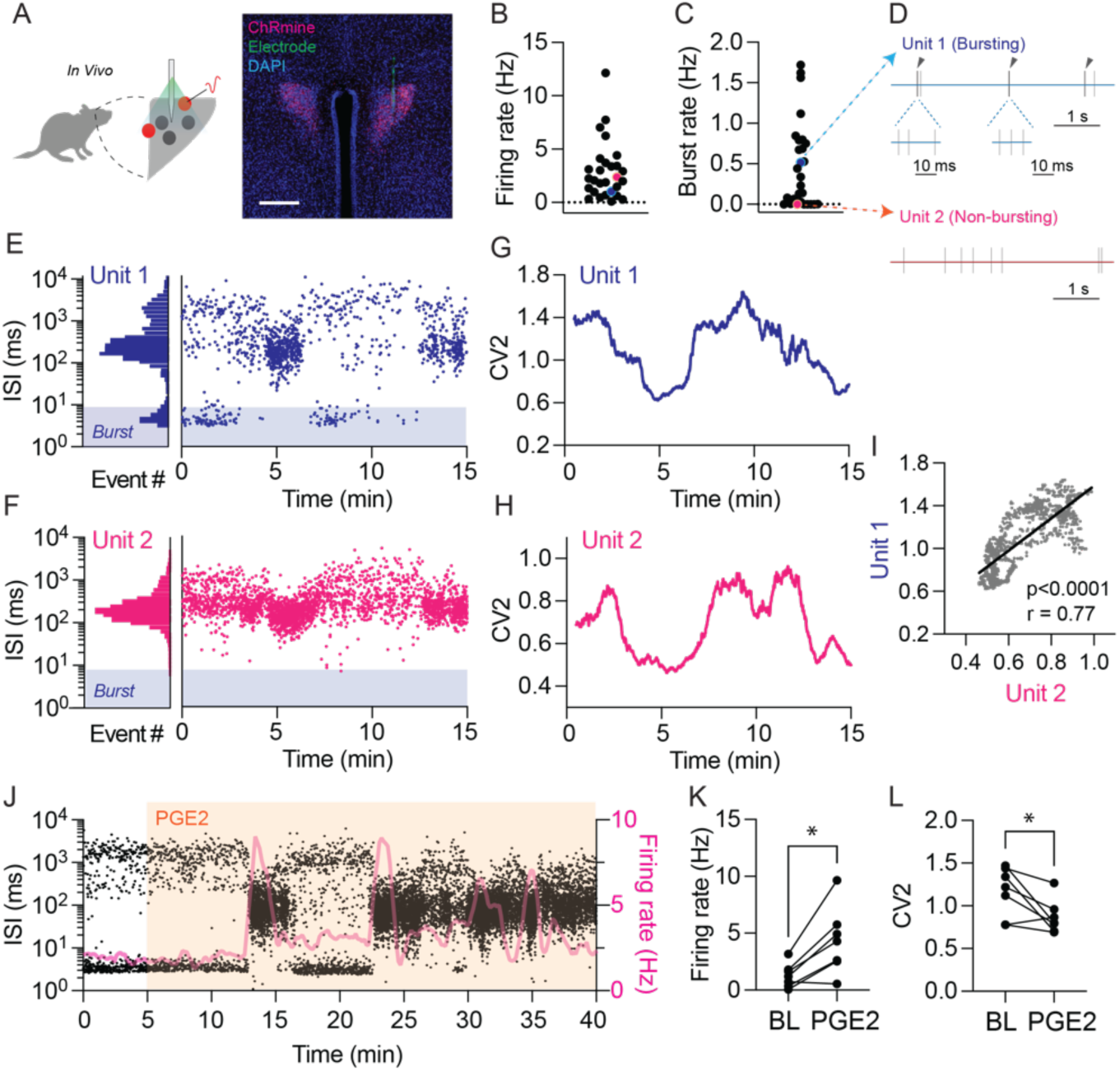
Single-neurons spiking patterns of CRHPVN neurons reflects internal states. **A**, *In vivo* recording and opto-tagging of CRHPVN neurons (left) with representative post-hoc validation of ChRmine expression and electrode placement (right). **B, C**, Summary of average firing rate (B) and burst rate (C). n = 28 from 18 mice. **D**, Example of simultaneously recorded busting neuron (top) and non-bursting neurons (bottom). **E, F**, Interspike interval (ISI) histogram (left) and ISI timecourse (right) for the bursting (E) and non-bursting (F) neurons. **G, H**, CV2 time courses for example neurons shown in E and F. **I,** Correlation of CV2 between two simultaneously recorded neurons. r = 0.77, p < 0.0001 **J,** Example ISI and firing rate time course before and after PGE2 administration (10 ng, icv). **K, L,** Summary of firing rate (K) and CV2 (L) changes after PGE2. Scale bar 50 μm, \**p* < 0.05

**Extended Data Fig. 2.**
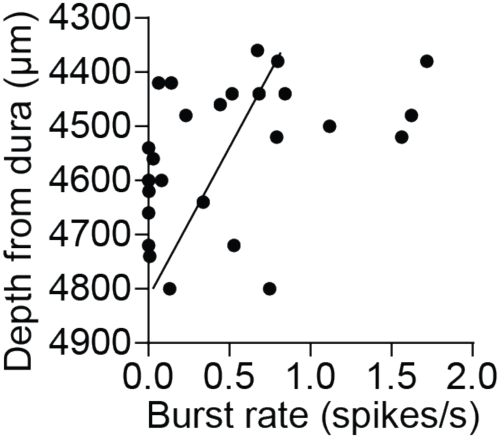
Relationship between burst rate and neuronal depth. Burst rate was significantly correlated with anatomical depth (P = 0.0155, r = -0.45).

Next, we examined stress-induced changes in functional states. To this end, we used prostaglandin E2 (PGE2), an inflammatory mediator, that excites CRH_PVN_ neurons and activates the HPA axis^15,16^. As expected, PGE2 (10 ng, icv) potently increased firing rate of CRH_PVN_ neurons, serving as a robust stress stimulus applicable to anesthetized animals. Importantly, the firing increase involved a clear shift in firing patterns, from rhythmic to tonic activity (Fig. 3J). Summary time course analysis revealed a long-lasting firing rate increase (Fig. 3K), which was accompanied by a decrease in CV2 (i.e., increase in regularity, Fig. 3L). These results demonstrate that a transition of functional states of CRH_PVN_ neurons, reflected in CV2 decrease, underlie stress-induced activity increase of CRH_PVN_ neurons. Built on these characterizations, we set out to test the causal roles of recurrent inhibitory circuit in regulating internal states using spiking patterns of CRH_PVN_ neurons.

### GABAergic inputs bidirectionally regulates CRH_PVN_ neurons functional states

Our model hypothesizes that recurrent inhibitory circuit bidirectionally regulates the low- and high-activity states of CRH_PVN_ neurons (Fig. 4A). This hypothesis generates explicit predictions for experimental testing. The first prediction is that the loss of GABAergic inhibition (i.e. disinhibition) increases the overall firing rate through tonic firing (i.e. low CV2). Conversely, the second prediction is that the increase of GABAergic inhibition decreases the firing rate through the emergence of rhythmic and irregular firing patterns (i.e. high CV2). To test these predictions, we chemogenetically inhibited or stimulated GABAergic inputs to CRH_PVN_ neurons. To this end, we crossed CRH-IRES-cre (Jackson Laboratory strain #012704) with vGAT-IRES-flp (Jackson Laboratory strain #029591). We injected flp-dependent AAVs into the PVN to retrogradely express DREADDs (hM4D(Gi) or hM3D(Gq)) in PVN projecting GABAergic (^PVN→^GABA) neurons (Fig. 4B). Hypothalamic areas immediately surrounding the PVN, including the perifornical area, anterior hypothalamus, zona incerta, and dorsomedial hypothalamus were robustly labelled (Fig. 4B), consistent with earlier studies identifying these PVN-surrounding areas as the major sources of GABAergic inputs to CRH_PVN_ neurons^17–19^.

**Fig. 4.**
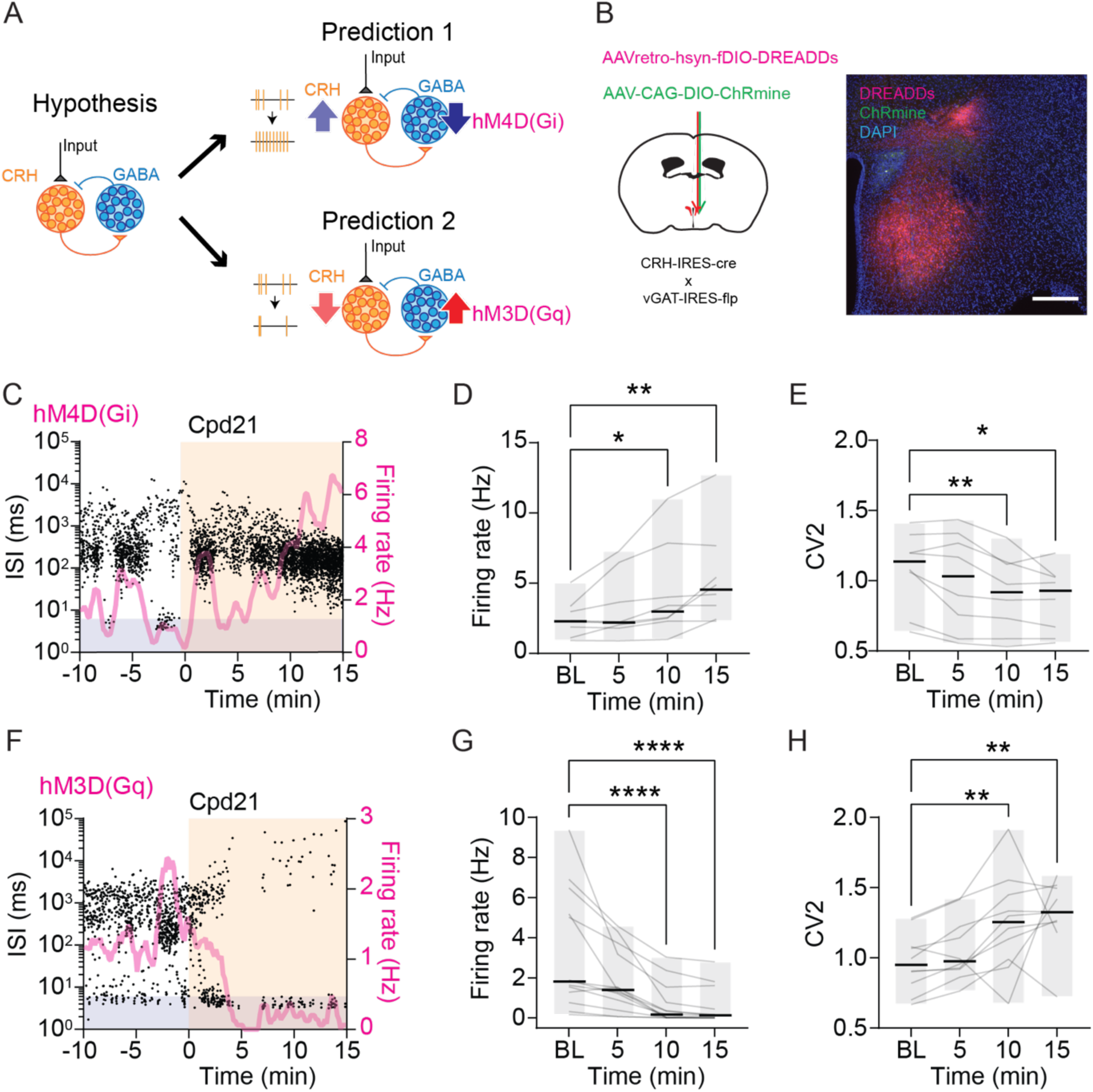
GABAergic afferents bidirectionally regulates CRHPVN neurons activity states. **A**, Recurrent inhibitory circuit model and predictions. **B.** AAV-mediated expression of ChRmine and DREADDs (left) with representative image of ChRmine and DREADDs expression (right). **C.** Example time course of interspike interval (ISI) and firing rate before and after chemogenetic inhibition of ^PVN→^GABAergic neurons. **D, E.** Summary time course of firing rate (D) and CV2 (E).n = 8 from 4 mice. **F.** Example ISI and firing rate time course before and after chemogenetic excitation of ^PVN→^GABAergic neurons. **G, H.** Summary time courses of firing rate (G) and CV2 (H). n = 13 (G) and n = 10 (H) from 8 mice. Scale bar 300 μm, \**p* < 0.05, ** *p* < 0.01, **** *p* < 0.0001

As predicted, chemogenetic inhibition of hM4D(Gi)-expressing ^PVN→^GABA neurons using the DREADD agonist Compound21 (Cpd21, 3 mg/kg i.p.) led to an increase in firing rate, accompanied by a shift in parring patterns from rhythmic to tonic (Fig. 4C). Time course analysis revealed a significant time-dependent increase in firing rate (Fig. 4C, D), with statistically significant elevations observed at 10 min and 15 min post-injection. This increase was accompanied by a reduction in CV2 (Fig. 3C, E), with significant decrease at 10 min and 15 min post-injection.

Conversely, chemogenetic activation of hM3D(Gq)-expressing GABAergic neurons suppressed firing rate of CRH_PVN_ neurons and induced a shift in firing patterns toward more rhythmic and irregular activity (Fig. 4F, G). Summary time course analysis showed a significant time-dependent decrease in firing rate (Fig. 4F-H), with significant reductions at 10 min (*Z* = -4.254, *p* < 0.0001) and 15 min (*Z* = -5.317, *p* < 0.0001) post-injection. Three out of thirteen recorded neurons were excluded from CV2 analysis due to complete silencing (Extended Data Fig. 2). Among the remaining neurons, a significant increase in CV2 was observed (Fig. 3F, G, I), with significant increases at 10 min and 15 min post-injection, corresponding to the time points of firing suppression.

Together, these results support prediction #1 and #2, demonstrating that a GABA-mediated network bidirectionally controls the functional states of CRH_PVN_ neurons.

**Extended Data Fig. 3.**
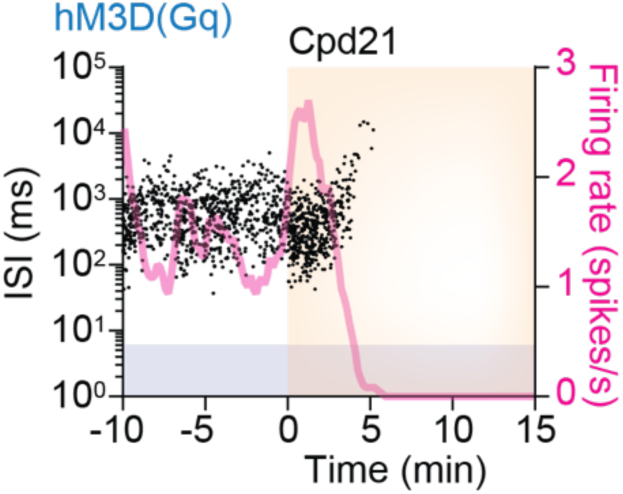
Chemogenetic excitation of PVN→GABAergic neurons silenced a subset CRHPVN neurons. Example time course of interspike interval (ISI) and firing rate before and after chemogenetic excitation of PVN→GABAergic neurons.

### CRH_PVN_ neurons drive feedback inhibition

The third prediction is that CRH_PVN_ neurons firings drive feedback inhibition (Fig. 5A). To test this, we optogenetically excited ChRmine-expressing CRH_PVN_ neurons and examined their firing activity. A 5 ms light pulse triggered an expected, time-rocked firing of CRH_PVN_ neurons, and this excitation was followed by a transient suppression of firing, most prominent around 200 ms post-stimulation (Fig. 5B-D). Summary time course analysis showed that firing rate decreased between 100 and 300 ms after the onset of light (Fig. 5C, D). This feedback inhibition supports our prediction.

**Fig. 5.**
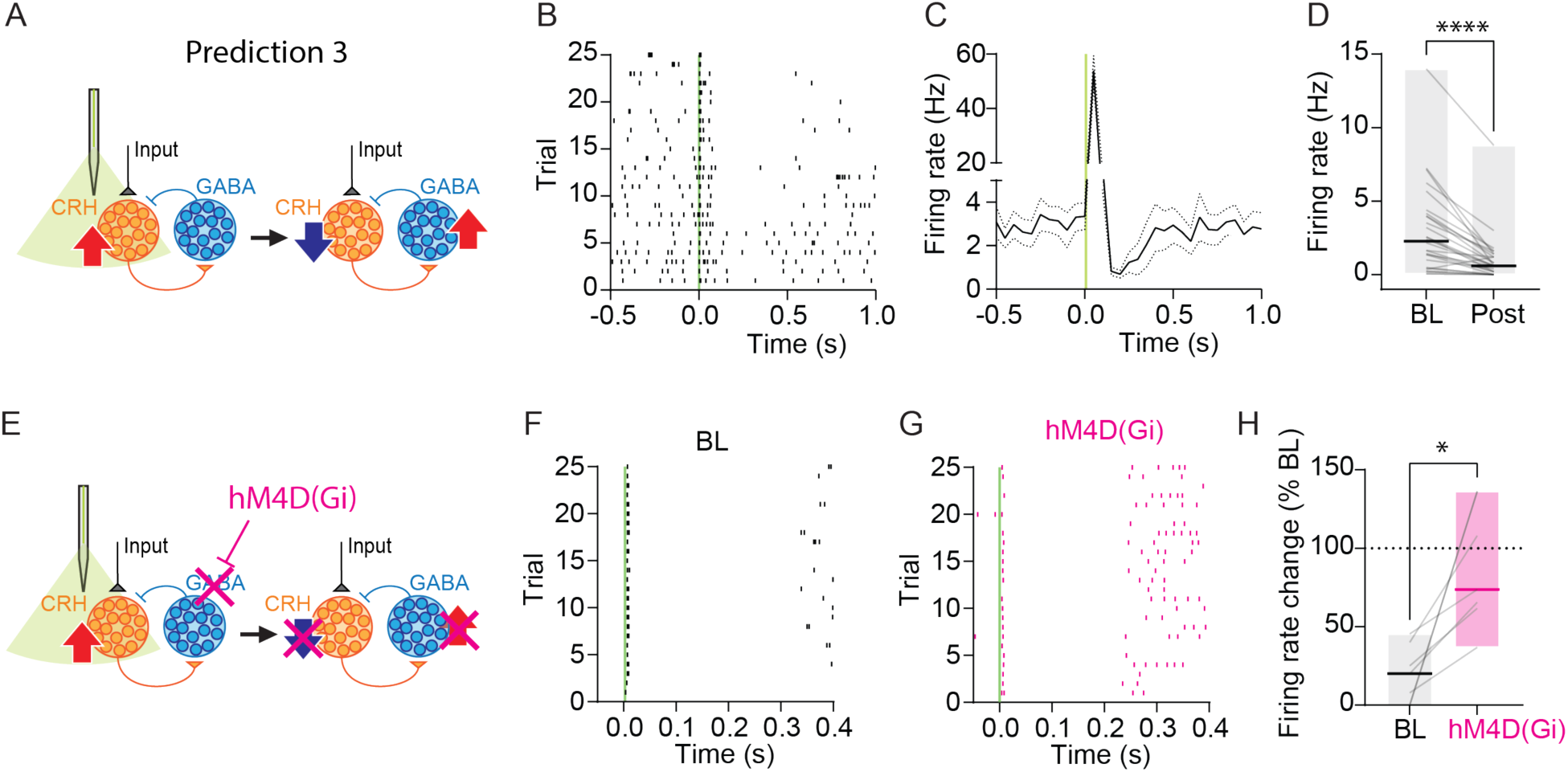
CRHPVN neurons drive GABAergic feedback inhibition. **A**, Model prediction. **B.** Example raster plot of CRHPVN neuron firing alighted to optogenetic stimulation (5 ms, green bar). **C.** Summary time course of firing rate before and after optogenetic stimulation. **D.** Firing rate before (BL) and after (Post, 100-300 ms) optogenetic stimulation. *p* < 0.0001, n = 28 from 18 mice. **E.** Schematic of chemogenetic inhibition of PVN→GABAergic neurons combined with optogenetic excitation of CRHPVN neurons. **F, G.** Example raster plots of CRHPVN neurons before (F) and after (G) chemogenetic inhibition of PVN→GABAergic neurons. **H.** Summary of optogenetically-induced feedback inhibition before and after chemogenetic inhibition of PVN→GABAergic neurons. *p* = 0.0156, N = 7 from 6 mice. \**p* < 0.05, \*\*\*\**p* < 0.0001

Next, we examined whether this light-induced feedback inhibition is indeed mediated by ^PVN→^GABA neurons. To this end, we compared the magnitude of inhibition before and after chemogenetic inhibition (via hM4D(Gi)) of ^PVN→^GABA neurons (Fig. 5E). Supporting our prediction, the chemogenetic inhibition reduced the magnitude of the light-induced feedback inhibition of CRH_PVN_ neurons (Fig. 5F-H). These results support our hypothesis that CRH_PVN_ neurons form recurrent inhibitory circuit that constrains their activity. Notably, the slow temporal scale (>100 ms) of the feedback inhibition is consistent with long ISIs that characterize the low-activity states (Fig. 2D, E) and point to that the recurrent inhibition involves slow neuromodulatory signaling.

### CRH receptors are dispensable for recurrent inhibition

Our model hypothesizes that CRH_PVN_ neurons release excitatory neurotransmitters to drive recurrent inhibition and constrain their activity (Fig. 3A). Previous experimental studies showed that CRH_PVN_ neurons induce slow excitation of their postsynaptic neurons via CRH-CRHR1 signalling^20–22^. Accordingly, our original computational model incorporates slow, CRHergic excitatory postsynaptic potentials to ^PVN→^GABA neurons ^12^. Together, the fourth prediction is that excitatory CRHergic transmission, via feedback inhibition, constrains the activity states of CRH_PVN_ neurons at low levels (Fig. 6A). To test this, we administered the CRH receptor antagonist Astressin (1 μg, icv) and examined firing activity of CRH_PVN_ neurons. Contrary to our prediction, CRH receptor blockade did not lead to a significant increase in CRH_PVN_ neurons firing rate (Fig. 6B, C) or significantly alter the firing variability (Fig. 6B). It should be noted, however, that the effects of Astressin were variable across units, with a subset exhibiting delayed excitatory responses (see around 15 mins in Fig. 6B). Astressin also did not have a significant effect on light-induced inhibition (Fig. 6E). Together, these results demonstrate that CRH receptors are dispensable for recurrent inhibition, at least in our experimental conditions, pointing to major roles for other excitatory neurotransmitters.

**Fig. 6.**
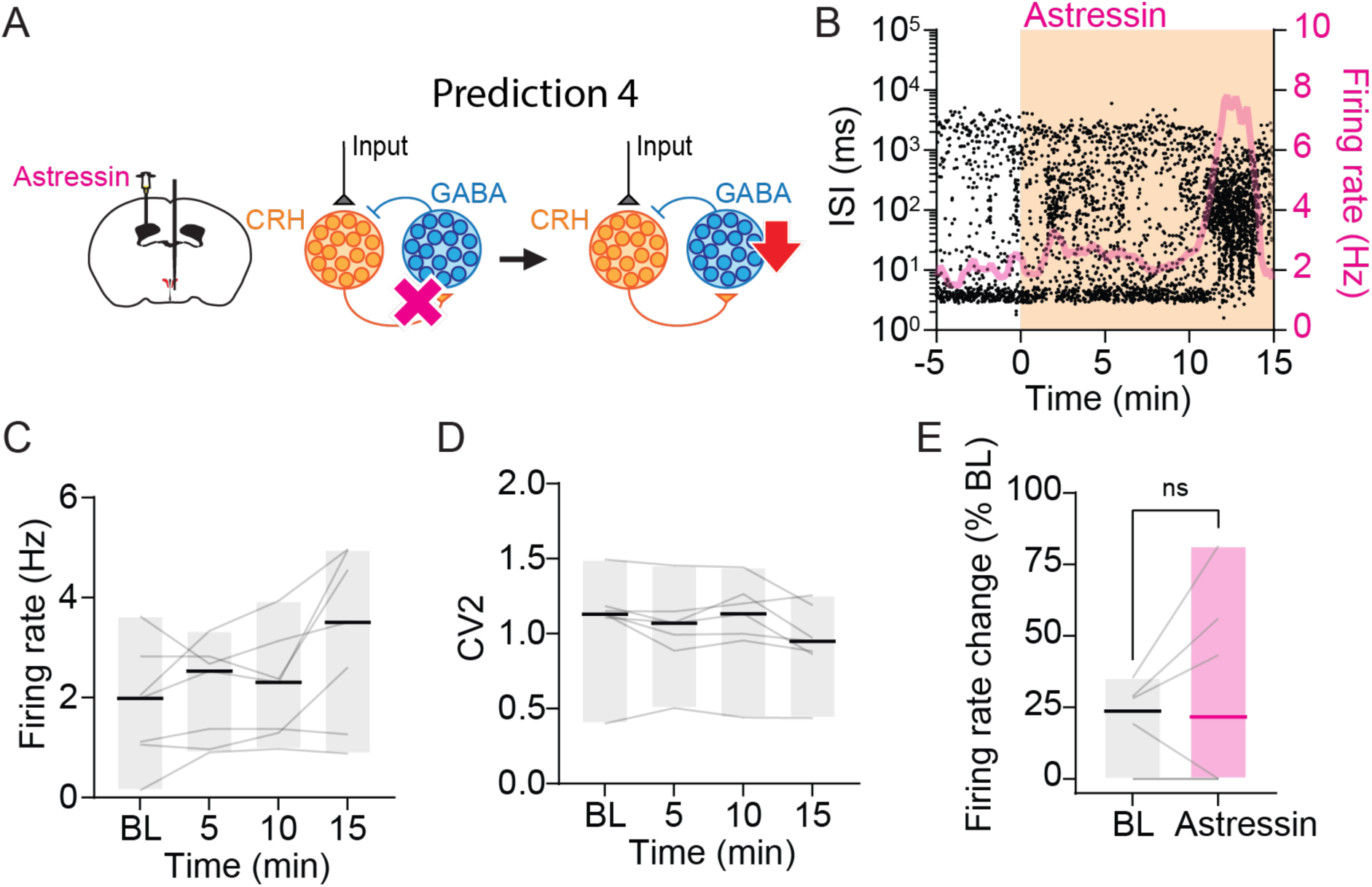
CRH receptor signaling is not required for recurrent inhibition. **A**, Model prediction. **B.** Example time course of interspike interval (ISI) and firing rate before and after injection of CRH receptor antagonist Astressin (1 μg, icv). **C, D.** Summary time course of firing rate (C) and CV2 (D). *p* = 0.016 (C) and *p* = 0.08 (D), n = 7 from 5 mice. **E.** Summary of optogenetically-induced feedback inhibition before and after chemogenetic inhibition of PVN→GABAergic neurons. *p* = 0.37, n = 6 from 6 mice.

### CRH_PVN_ neurons forms monosynaptic glutamatergic transmission onto ^PVN→^GABA neurons

The majorities, if not all, CRH_PVN_ neurons express vGlut2^23–25^ and release glutamate onto some of their postsynaptic targets^3^. To investigate whether CRH_PVN_ neurons release glutamate onto ^PVN→^GABA neurons, we used *ex vivo* patch-clamp electrophysiology in acute slices prepared from *CRH-IRES-cre x vGAT-IRES-flp* mice where ^PVN→^GABA neurons were retrogradely labelled with mCherry for visual identification, and CRH_PVN_ neurons expressed ChRmine-eYFP for optogenetic stimulations (Fig. 7A). Under the voltage-clamp recording, brief light illuminations (20 ms) reliably triggered rapid postsynaptic currents in a subset of ^PVN→^GABA neurons (46%, 51/110 neurons, Fig. 7B-D). Bath application of AMPA/kainite receptor antagonist DNQX (10 μM) abolished the optogenetically-evoked postsynaptic currents (oPSCs, Fig. 7B), validating glutamatergic transmission (Fig. 7B, C inset). Furthermore, bath application of TTX (1 μM) abolished the oPSCs while subsequent application of 4AP (40 μM) restored them (Fig. 7B, C), indicating that they are monosynaptic connections^26^. The oPSC-positive neurons were distributed in areas surrounding the PVN, including the perifornical area, lateral hypothalamus, anterior hypothalamus and zona inserta (Fig. 7D). Post-recording histology of ^PVN→^GABA neurons revealed close appositions between their dendrites with ChRmine-eYFP expressing fibers originating from CRH_PVN_ neurons (Fig. 7E). Together, these results demonstrated that CRH_PVN_ neurons form monosynaptic glutamatergic synapses onto ^PVN→^GABA neurons.

**Fig. 7.**
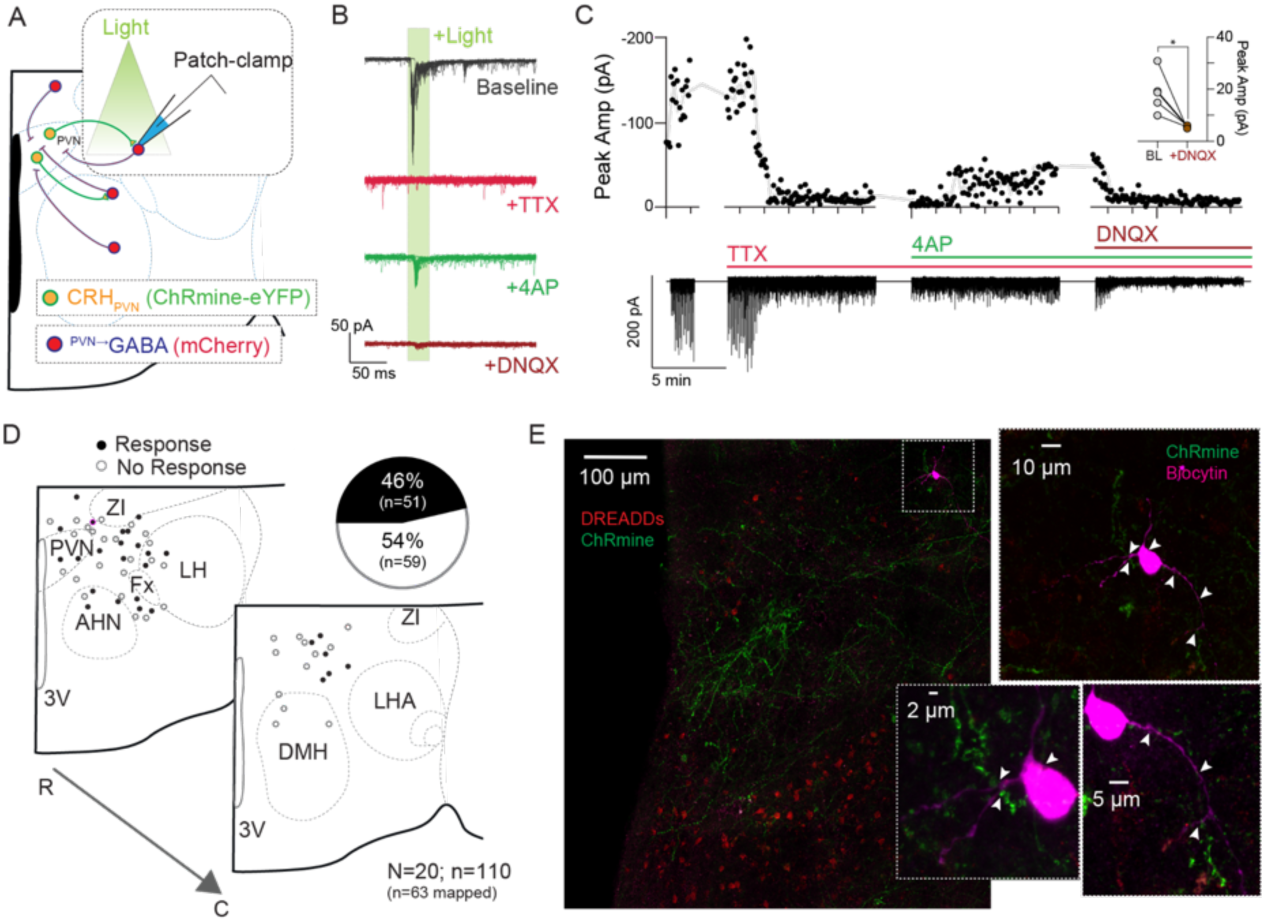
CRHPVN neurons form monosynaptic glutamatergic connections onto PVN→GABA neurons. **A**, Experimental setup: optogenetic stimulation of CRHPVN efferents (green) with patch-clamp electrophysiology recordings from PVN→GABAergic neurons (red). **B.** Example traces of optogenetically-evoked postsynaptic currents (oPSCs) before and after drug application. **C.** Time course of oPSCs amplitude (top) and corresponding raw trace (bottom): inset shows summary of amplitude reduction by AMPA/kainite receptor antagonist DNQX. **D.** Distribution of PVN→GABA neurons responding with (filled circle) or without (empty circle) oPSCs. **E.** Biocytin-filled PVN→GABA neuron (purple). Insets show higher magnification images with ChRmine-eYFP labelled fibers in close apposition to dendrites of the recorded, biocytin filled neuron (arrowheads). PVN→GABA neurons express mCherry (red) and distributed around the PVN.

### Data-driven refinements of recurrent inhibition models

Informed by these new experimental results suggesting that CRH_PVN_ neurons drive the recurrent inhibition via fast, glutamatergic excitation, we refined our computational modeling. The original model (Fig. 8A) incorporated slow excitatory transmission (τ*_e(_*_CRH→GABA)_ ≍ 200 ms) in CRH→GABA synapses, and fast inhibitory transmission (τ*_i(GABA_*_→_*_CRH)_* ≍ 20 ms) in GABA→CRH synapses (E*_slow_*/I*_fast_*, Fig. 8A)^12^. For the refinement, we replaced the slow CRH→GABA excitation with fast τ*_e(_*_CRH→GABA)_ (≍ 10 ms) and paired with either the original fast inhibition (E*_fast_*/I*_fast,_* Fig. 8B) or slow inhibition (τ*_i(GABA_*_→_*_CRH)_* ≍ 30 ms, E*_fast_*/I*_slow_*, Fig. 8C). Notably, both refined models, along with the original E*_slow_*/I*_fast_* model, reproduced the key *in vivo* features: inhibition-constrained spiking dynamics of CRH_PVN_ neurons at baseline and disinhibition-induced transitions to tonic spiking (Fig. 8A-E bottom). More generally, CRH_PVN_ neurons spiking dynamics could be captured across a broad range of τ*_e(_*_CRH→GABA)_ and τ*_i(GABA_*_→_*_CRH)_* combinations (Fig. 8F), with simulations systematically evaluated by comparing log ISI distributions of simulated and representative *in vivo* spike trains using mean Wasserstein Distance (see Materials and Methods).

**Fig. 8.**
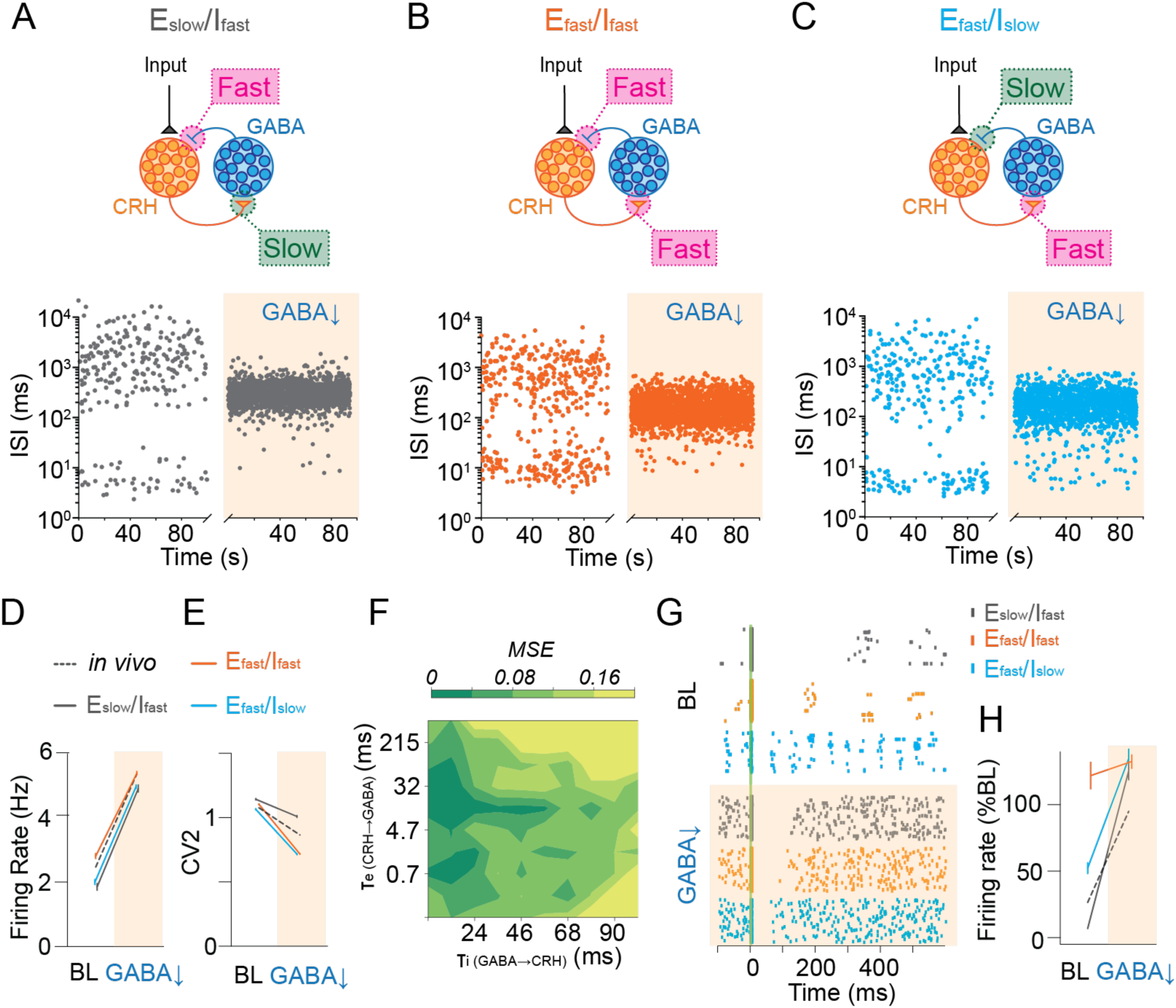
Computational model refinements. **A**, Original model with slow CRH→GABA excitation and fast GABA→CRH inhibition (top). Example interspike interval (ISI) time couse at baseline and after disinhibition (bottom). **B.** Model with fast CRH→GABA excitation and fast GABA→CRH inhibition (top) and corresponding ISI time course (bottom). **C.** Model with fast CRH→GABA excitation and slow GABA→CRH inhibition (top) and corresponding ISI time course (bottom). **D, E,** Summary plots of firing rate and CV2 for three models compared with *in vivo* data adopted from Fig. 4D, E. **F.** Heatmap of mean squared error (MSE) across combinations of excitatory CRH→GABA and inhibitory GABA→CRH synaptic time constants (τ). **G.** Example raster plots showing optogenetically-induced feedback inhibition in three models at baseline and after disinhibition. Vertical green bar indicates time-locked excitation of model CRH neurons mimicking optogenetic excitation *in vivo*. **H.** Summary plots of light-induced feedback inhibition across three models and in vivo data adopted from Fig. 5H at baseline and after disinhibition. Error bars indicate SEM.

We next examined which models could also account for the protracted feedback inhibition of CRH_PVN_ neurons observed after optogenetic stimulation *in vivo* (Fig. 5). This test revealed that the both the E*_slow_*/I*_fast_* and E*_fast_*/I*_slow_* models, but not the E*_fast_*/I*_fast_* model, reproduced the *in vivo*-like sustained inhibition (Fig. 8G, H), underscoreing the importance of slow synaptic time constants in recurrent inhibition. Consistent with *in vivo* experiments, dampening GABA→CRH inhibitory transmission (mimicking chemogenetic inhibition of ^PVN→^GABA neurons) effectively abolished simulated feedback inhibition, validating the requirement of recurrent inhibition motif for the feedback inhibition. (Fig. 8G, H). A recent study identified a feedforward inhibitory circuit targeting CRH_PVN_ neurons^27^, a circuit motif known to control state transitions in other brain areas^28^. We therefore additionally tested a feedforward inhibition model (Extended Data Fig. 4). While feedforward inhibition could reproduce transitions in spiking dynamics, it failed to account for the protonated feedback inhibition (Extended Data Fig. 4). These results highlight recurrent inhibition as a critical circuit motif for regulating CRH_PVN_ neurons spiking dynamics *in vivo*.

**Fig. 8.**
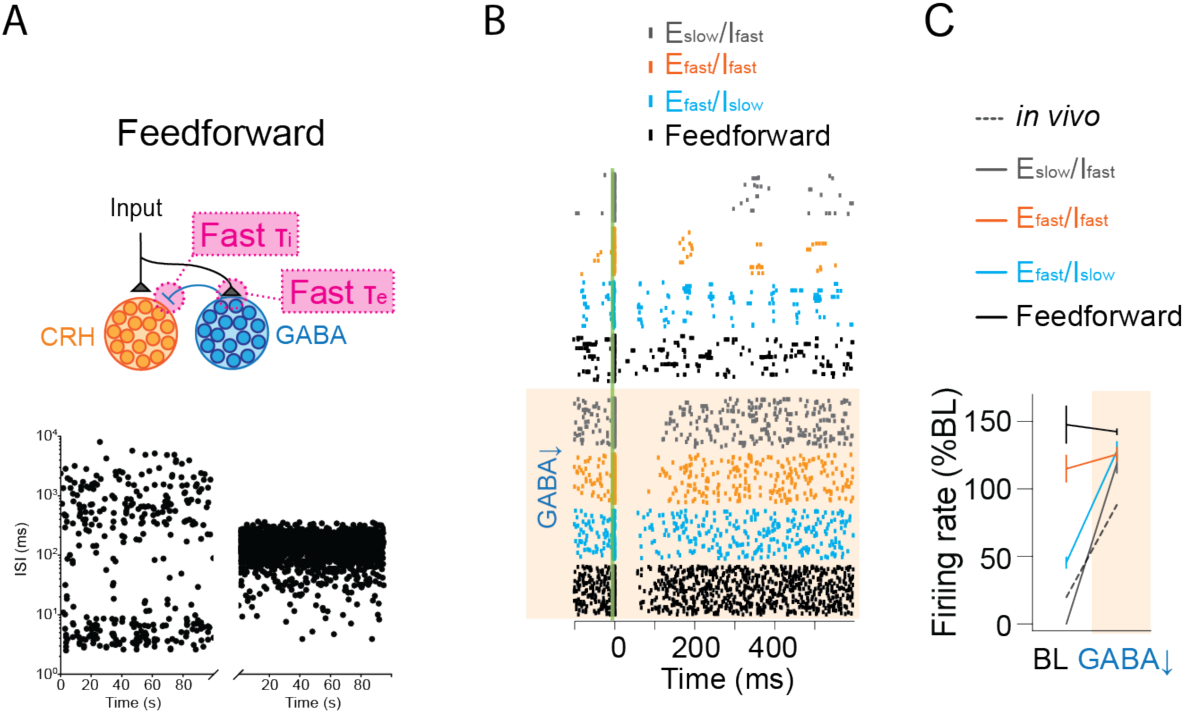
Feedforward inhibition model. **A**, Model with feedforward inhibiton with fast excitatory inputs to GABA neurons and fast inhibitory inputs to CRH neurons (top). Example interspike interval (ISI) time couse at baseline and after disinhibition (bottom). **B.** Example raster plots showing optogenetically-induced feedback inhibition in four models at baseline and after disinhibition. Vertical green bar indicates time-locked excitation of model CRH neurons mimicking optogenetic excitation *in vivo*. **C.** Summary plots of light-induced feedback inhibition across four models and in vivo data adopted from Fig. 5H at baseline and after disinhibition. Error bars indicate SEM.

## DISCUSSION

Here, we provide evidence for a novel hypothalamic recurrent circuit that regulates functional states of CRH_PVN_ neurons. Our findings fundamentally revise the prevailing circuit framework, shifting from the long-held view that converging inputs to CRH_PVN_ neurons represent a one-way flow of signals driving the stress response, to one in which these neurons are embedded in recurrent connectivity that constantly shapes their functional states. Specifically, under non-stress conditions, recurrent inhibitory circuits provide crucial feedback inhibition, constraining these neurons in low-activity states, whereas, in response to stress, a loss of recurrent inhibition permits rapid transitions to high-activity states. This circuit mechanism was generalizable across different modalities of stress, including novel environment (Fig. 1, 2), inflammatory stress (Fig. 3), and pain-related nerve stimuli^12^, endowing key bidirectional control over CRH_PVN_ neurons activity. Additionally, our study highlights the power of iterative experiment–model integration where *in vivo* neural dynamics data led to computational circuit model, and the model subsequently provided experimentally testable hypotheses for biological circuit structures and operations, and lastly, model-guided experiments in turn revealed key model refinements and generate new predictions.

Decades of work have established that GABAergic inputs play key regulatory roles in CRH_PVN_ neurons, with more than half of synaptic terminals apposing to the soma and dendrites of CRH_PVN_ neurons^29^. Loss of GABA_A_ receptor-mediated inhibition (i.e. disinhibition), whether by pharmacological blockade^30^ or stress^31–33^, strongly excites CRH_PVN_ neurons and activates downstream hormonal responses. Our finding of recurrent inhibition aligns with these established mechanisms but provides an important revision: a subset of these GABAergic inputs forms recurrent connections with CRH_PVN_ neurons, creating a feedback inhibitory loop. To our best knowledge, this study is the first to report *in vivo* single unit activities of CRH_PVN_ neurons during their optogenetic stimulation, and their spiking patterns provided direct evidence for this feedback inhibition, which was attenuated by chemogenetic inhibition of ^PVN→^GABAergic neurons. This recurrent inhibitory motif provides circuit-level explanations for earlier observations that glutamate microinjection into the PVN produced little excitation of CRH_PVN_ neurons^30^, while tonic glutamatergic drive is nonetheless required for disinhibition-induced excitation^34^. Together, these results support a model in which recurrent inhibition constrains the gain of CRH_PVN_ neurons^12^, crucial for stabilizing their baseline activity.

In response to stress, CRH_PVN_ neurons rapidly and reversibly transition to high-activity states, suggesting that stress disengages recurrent inhibitory circuits. Likely sources of such modulatory inputs are stress-responsive corticolimbic regions, including the prefrontal cortex, hippocampus and amygdala, that form classic disinhibitory pathways through dense axonal projections to peri-PVN regions populated by ^PVN→^GABAergic neurons^35–38^. Additionally, recent studies have revealed axo-axonic modulations of GABAergic terminals onto CRH_PVN_ neurons via presynaptic GABA_B_ receptors^39^, which plays key roles in HPA axis activation during fasting^33^. Further, inflammatory mediator PGE2 dampens GABA release onto CRH_PVN_ neurons via presynaptic EP3 receptors^16^. Together, these findings highlight diverse mechanisms by which recurrent inhibition can be modulated, providing multiple paths through which distinct stressors converge to influence the activity states of CRH_PVN_ neurons.

Contrary to our original model prediction, CRHR antagonist had little effect on baseline low-activity firing dynamics, indicating that CRHR signalling is dispensable for recurrent inhibition. Our original model^12^ incorporated slow excitatory transmission (τ*_e(_*_CRH→GABA)_ ≍ 200 ms) because 1) rhythmic protonated silence features the low-activity state and 2) a subset of CRHR1-expressing neurons within and around the PVN are GABAergic and form recurrent circuits with PVN_CRH_ neurons^21,22,40^. To refine the model, we experimentally validated that CRH_PVN_ neurons, which express vGlut2^23–25^, form glutamatergic excitatory synapses onto ^PVN→^GABAergic neurons, consistent with a prior report^3^. Parameter scans of the updated models revealed that a broad range of both excitatory (τ*_e(_*_CRH→GABA)_) and inhibitory (τ*_i(_*_GABA→CRH)_) synaptic time constant can replicate functional state transitions observed *in vivo*. Notably, however, the new model identified slow inhibitory (τ*_i(_*_GABA→CRH)_) kinetics, along with the original slow excitation (τ*_e(_*_CRH→GABA)_), as critical for sustaining the optogenetically-induced, protracted feedback inhibition, thereby generating a new prediction for the biological mechanisms of ^PVN→^GABAergic→CRHPVN transmission contributing to feedback inhibition.

In summary, we provide converging biological and computational evidence that hypothalamic recurrent inhibition regulates stress-driven activity transitions of CRH_PVN_ neurons. More broadly, our work highlights the value of iterative integration between modeling and experiment to uncover circuit mechanisms governing the dynamic regulation of stress responses.

## METHODS

### Mice

All animal protocols were approved by the Western University or the University of Calgary Animal Care and Use Committee. For the miniature microscopy experiments, the heterozygous offsprings of Crh-IRES-Cre (B6(Cg)-Crhtm1(cre)Zjh/J; stock number 012704) mice crossed with Ai148 (Ai148(TIT2L-GC6f-ICL-tTA2)-D; stock number 030328) mice were utilized. For *in vivo* electrophysiology experiments, the heterozygous offsprings from Crh-IRES-Cre mice crossed with Ai14 (B6.Cg-Gt(ROSA)26Sortm14(CAG-TdTomato)Hze/J; stock number 007914) or with vGAT-Flp ( B6.Cg-Slc32a1tm1.1(flpo)Hze/J; stock number 029591) mice were used. Breeder mice were obtained from Jackson Laboratories.

Mice were housed on a 12-h:12-h light:dark cycle (lights on at 7:00 a.m.) with ad libitum access to food and water. All subjects were randomly assigned to different experimental conditions used in this study. Mice were 6-10 weeks old at the time of surgery and virus injection.

### Stereotaxic GRIN lens implantation for miniature microscopy

Mice were maintained under isoflurane anesthesia in the stereotaxic apparatus. The GRIN lens (6.1 mm length; Inscopix) was lowered at a 100 µm/min speed using a motorized stereotaxic apparatus. Implantations were targeted to dorsal to the PVN (A/P: −0.7 mm, M/L: 0.2 mm, D/V: −4.2 mm from the dura) of CRH-IRES-Cre: Ai 148 mice and were affixed to the skull with METABOND® and dental cement. At least one month after lens implantation, a baseplate was installed on the head. Experiments started after an additional two weeks of recovery and handling.

### Stereotaxic AAV microinjection

Mice were maintained under isoflurane anesthesia in the stereotaxic apparatus. AAV injections were performed using Nanoject II (Drummond Scientific) at a rate of 2 nL/s targeted to the PVN bilaterally (A/P: −0.7 mm, M/L: ±0.25 mm, D/V: −4.7 mm from the dura). After each injection, the pipette was left in place for 5 minutes to allow diffusion before being slowly withdrawn. At the end of surgery, mice received subcutaneous fluid therapy (saline, 10 mL/kg) and an analgesic (carprofen, 5 mg/kg). To express ChRmine in CRH_PVN_ neurons, a Cre-dependent AAV (AAV2/9-CAG-DIO-ChRmine-EYFP; RRID: SCR_016477; Canadian Neurophotonics Platform Viral Vector Core Facility) was injected into CRH-IRES-Cre; Ai14 mice. 25-50 nL (3 x 10^12^ GC/mL) was injected into the PVN. To express both ChRmine in CRH_PVN_ neurons and DREADDs in ^PVN→^GABAerginc neurons, CRH-IRES-Cre; vGAT-IRES-Flp mice were first injected with a retrograde AAV that express DREADDs (hM3D(Gq)-mCherry or hM4D(Gi)-mCherry) in a Flp-dependent manner into the PVN. Approximately two weeks after the injection, second surgery was performed to inject a Cre-dependent AAV that express ChRmine into the PVN. The following AAV_retro_ were used (AAV_retro_-hsyn-fDIO-hM3D(Gq)-mCherry-wPREpA, Addgene # 154868-AAVrg; or AAV_retro_-hSyn-fDIO-hM4D(Gi)-mCherry-WPREpA, Addgene #154867-AAVrg). 60 nL of AAV_retro_ (7 x 10^12^ vg/mL) was injected into the PVN. Experiments started after at least additional two weeks of recovery and handling.

### Miniature microscopy data analysis

Activity of CRH_PVN_ neurons was recorded continuously for at least 15 minutes at 20 FPS, 40 % LED power and 2.5 gain, using the nVista Acquisition Software (Inscopix). The video files were cropped with ImageJ and the data analysis was performed by the Min1pipe script^41^ in MATLAB. Peaks of spontaneous activity were defined as local maximum points above a threshold calculated as the proportion of the mean absolute deviation plus the mean of the low-pass filtered df/f, on a cell to cell basis:

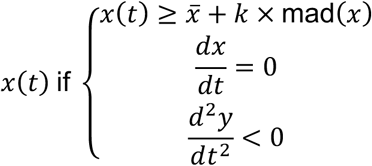

Where *x* is 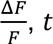 is time, mad is the mean absolute deviation, *k* was set to 2, and the cut-off frequency was set to 5Hz.

### Electrophysiology

Animals were initially anesthetized under isoflurane (1–2%) and urethane (1.5 g/kg in 0.9% saline, intraperitoneal) to perform craniotomy for insertion of the recording probe into the brain. The Neuropixels 1.0 rodent probe (IMEC) was mounted to a stereotaxic arm (Kopf Instruments) and inserted vertically into the brain toward the PVN (A/P: –0.70 mm, M/L: ±0.25 mm). The probe was slowly lowered past the PVN to a depth of –5.40 mm from the dura, with the window for finding PVN neurons defined to be between –4.35 mm and –4.85 mm from the dura. The electrode was allowed to settle for at least 45 minutes to minimize recording drift before beginning baseline recordings. Additionally, an optic fiber (see details in *Optogenetic Activation of CRH Neurons*) was mounted to a separate stereotaxic arm and inserted at a 12-degree angle from vertical, targeting a region approximately 300 μm dorsolateral to the PVN. The exposed portion of the probe shank and optic cable connection were coated in black paint to minimize light spill during optogenetic stimulation. For intracerebroventricular drug delivery experiments (see below), an additional stereotaxic arm was used to lower a fine glass capillary pipette into the lateral ventricle (A/P: –0.50 mm, M/L: 1.0 mm, and D/V: −2.0 mm from bregma). Following all placements, isoflurane anesthesia was discontinued, and the animal remained on urethane for the remainder of the recording session.

The Neuropixels 1.0 probe was connected to an IMEC headstage. Digitized signals were transmitted to the IMEC PXIe-8381 acquisition module, housed within a PXIe-1082 chassis (National Instruments). Data were streamed to the acquisition computer. All data acquisition was managed using SpikeGLX software, which enabled synchronized visualization and recording of electrophysiological signals. Recordings were sampled at 30 kHz per channel.

Spike sorting was performed offline using Kilosort v2.5. Clusters labeled “Good” by Kilosort— indicating high isolation and waveform consistency—were manually curated using Phy. Only well-isolated, validated units were included in further analyses.

### Optogenetics

Optogenetic stimulation was triggered by TTL pulses routed from the RZ5D processor (Tucker-Davis Technologies) to a PlexBright LD-1 Single Channel LED Driver (Plexon). The LED driver regulated the output of a PlexBright Table-Top LED Module (Lime, 550 nm), which was coupled to a 200 μm core optical patch cable (PlexBright, Plexon). The patch cable was connected via a plastic sleeve to an implanted optical fiber (200 μm core diameter, 0.66 NA, 2 mm active length; Optogenix, Lambda Fiber Cannulae) targeting the PVN. For optogenetic identification of CRH_PVN_ neurons, light pulses of 5 ms duration were delivered in 25 trials at 3 s intervals with a power output of 0.5 mW. To store stimulus timings synced to the neural recording, the TTL output from the RZ5D was split and connected to the SMA input on the Neuropixels basestation. These stimulus pulses were recorded on channel 385 in SpikeGLX, which captures external inputs. Stimulus onset times were later extracted by thresholding the recorded waveforms on this channel.

### DREADDs

In DREADD-expressing animals, DREADD agonist 21 dihydrochloride (C21; TOCRIS 6422) was delivered via intraperitoneal injection at 3 mg/kg dissolved in saline.

### Intracerebroventricular drug infusion

Astressin (Tocris Bioscence, Cat. No. 1606) was dissolved in saline. Mice were injected ICV using a gravity-pulled glass needle connected to a 20 uL Hamilton syringe filled with mineral oil. 1 µg of Astressin dissolved in 5 µL of sterile saline was pressure injected into the lateral ventricle at a rate of 2 µL/min. PGE2 (Cayman Chemical, Cat. No. 14010) was dissolved in ethanol at 1 mM concentration as a stock. The stock was diluted in saline at the concentration of 28 μM, and 1 μL was injected in the lateral ventricle at the rate of 2 μL/min.

### Histology

At the end of recording, animals were euthanized with an overdose of pentobarbital sodium (150 mg/kg, i.p.) and transcardially perfused with 0.9% saline, followed by 4% paraformaldehyde (PFA). The brain was postfixed in 4% PFA at 4°C overnight. The brain was washed in phosphate buffer saline (PBS) solution and sliced into 40 μm coronal sections and counter stained with 4,6-diamidino-2-phenylindole (DAPI). The slices were mounted and verified for CRH expression (tdTomato), ChRmine expression (eYFP), DREADDs expression (mCherry) and/or the dye-painted electrode tract (Vybrant DiD, Thermo Fisher V22889).

### Data Analysis and Statistics

Data analyses were carried out using built-in and custom-built software in MATLAB (MathWorks). Graphs were made and statistical analysis were conducted using Prism 10 (GraphPad). All pairwise and repeated measures comparisons were performed using the Wilcoxon signed-rank test or Friedman test, respectively, with statistical significance assessed at p < 0.05.

#### Optogenetic identification of CRH_PVN_ neurons

Following offline sorting of waveforms, ChRmine-expressing CRH single units were identified by their time-locked responses to light. Neurons were considered light-responsive if they showed a significant increase in spiking activity in response to a 5 ms pulse of 550 nm lime-colored light across 25 trials. Spike rates in the 10 ms windows immediately before and after light onset were compared using a paired t-test. A significant increase in post-stimulus firing was used as the primary criterion for light responsiveness. As a secondary criterion, the response probability within the 10 ms post-stimulus window was required to exceed 60%. Response probability was calculated as the proportion of trials in which at least one spike occurred within the defined time window. Only units that met both the statistical significance and response probability thresholds were classified as light-responsive and thus identified as CRH_PVN_ neurons.

#### Firing pattern analysis

Firing patterns were characterized by calculating interspike intervals (ISI), defined as the time between consecutive spikes. ISI time courses were plotted by aligning ISI value to the time of the preceding spike (i.e., the start of the interval). Bursts were detected as series of two or more spikes starting with the initial ISI less than 6 ms and subsequent ISIs less than 20 ms.

Firing variability was quantified using the CV2 metric, which estimates local variability in ISI by comparing adjacent ISIs rather than computing global variance. This makes CV2 less sensitive to slow changes in firing rate and better suited for analyzing dynamic states of neural activity (Holt et al., 1996). For a given pair of consecutive ISIs (ISI₁ and ISI₂), CV2 is defined as:

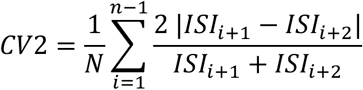

This measure approaches 1 for Poisson-like spiking and increases with irregularity. Neural states were classified as high or low variability based on CV2 distributions. CV2 correlation between units was assessed using Pearson correlation of average CV2 values computed within 30-second sliding windows (1-second step size).

#### Firing rate analysis

The average baseline firing rates were calculated as the total number of spikes divided by total time during the spontaneous baseline recording (10 min), or 5 minute analysis windows of each experiment. The firing rate time course was calculated as the number of spikes in every 10 s bins. The running average firing rate represents a moving average of five consecutive bins (bin[*i*-2]+bin[*i*-1]+bin[*i*]+bin[*i*+1]+bin[*i*+2]/5) and was used to visualize spontaneous fluctuations of firing rate. These fluctuations in firing rate were then overlaid onto ISIs plotted across time to visualize the relationship between firing rate to firing patterns.

#### Chemogenetic and pharmacological time course analysis

To assess the effects of chemogenetic and pharmacological interventions, CRH_PVN_ neuron firing rates and firing patterns were compared between a 5-minute pre-injection BL and successive 5-minute windows following drug administration, up to 15 minutes post-injection. All comparisons were tested by Friedman test with post-hoc Dunn’s multiple comparisons test.

#### Optogenetic feedback inhibition analysis

The time course of CRH_PVN_ optogenetic activation to a 5 ms light pulse was plotted in 50 ms bins, averaged over 25 trials within each unit and then across units. To assess firing rate change before and after optogenetic stimulations, the average firing rate during the 1-second pre-stimulation BL was compared to the firing rate between 100 and 300 ms after light onset. To evaluate the effects of DREADD(Gi) and Astressin on this feedback inhibition, the post-stimulation firing rate was expressed as a percentage of baseline (to control for baseline changes) and compared across conditions with and without these manipulations (Wilcoxon signed-rank test).

### Ex Vivo Electrophysiology

Mice were anesthetized with isoflurane and quickly decapitated. Brains were rapidly removed after decapitation and placed into a cutting solution containing the following (in mM): 87 NaCl, 2.5 KCl, 25 NaHCO3, 0.5 CaCl2, 7 MgCl2, 1.25 NaH2 PO4, 25 glucose and 75 sucrose (Osmolarity: 315–320 mOsm), saturated with 95% O2/5% CO2. Coronal sections (250 μm thick) containing the hypothalamus were cut using a vibratome (VT-1200, Leica Biosystems). The aCSF solution consisted of the following (in mM): 123 NaCl, 2.5 KCl, 1.25 NaH2PO4, 26 NaHCO3, 10 glucose, 2.5 CaCl2 and 1.5 MgCl2, saturated with 95% O2/5% CO2, pH 7.4, osmolarity 300 mOsm. Slices were recovered at 34 °C in artificial cerebrospinal fluid (aCSF) for 45 min and subsequently kept at room temperature. During experimentation, slices were perfused at a rate of 2 ml/min in aCSF and maintained at 27°C–30°C.

Borosilicate glass micropipettes (BF120-69-15, Sutter Instruments) were pulled in a Flaming/Brown Micropipette Puller (P-1000, Sutter Instruments) and filled with an intracellular fluid containing the following (in mM): 108 K-gluconate, 2 MgCl2, 8 Na-gluconate, 1 K2-ethylene glycol-bis(β-aminoethyl ether)-N,N,N′,N′ -tetraacetic acid (EGTA), 4 K2-ATP, 0.3 Na3-GTP, 10 HEPES (osmolarity: 283‒289 mOsm and pH: 7.2‒7.4). The resistance of the pipettes was between 3 and 5 MΩ. To enable the localization of patched neurons, a 0.5% (w/v) concentration of biocytin (B1592, Thermo-Fisher Scientific) was added to the internal solution on the day of the experiment.

For the characterization of intrinsic excitability, heterozygous offsprings from Crh-IRES-Cre mice crossed with Ai14 mice were used to enable visual identification of CRHPVN neurons in acute slices. Recordings were obtained in the presence of antagonists for both AMPA/Kainite receptors (20 μM DNQX) and GABAA receptors (100 μM picrotoxin).

For optogenetic examinations of synaptic transmission in CRH→^PVN→^GABA synapse, the heterozygous offsprings from Crh-IRES-Cre mice crossed with vGAT-Flp mice were used. To express both ChRmine in CRHPVN neurons and mCherry in ^PVN→^GABAerginc neurons, AAVs were injected as described above, and experiments were performed 3-4 weeks after the injection. To isolate EPSCs, picrotoxin (100 μM) was bath applied. EPSCs were recorded in voltage-clamp mode with the membrane voltage held at –70 mV. Optogenetically-evoked excitatory post-synaptic potentials (oEPSCs) were elicited with pairs of 20-ms pulses of 473 nm light, delivered with a 50-ms inter-pulse interval at 5-sec sweeps. To confirm monosynaptic connectivity, action-potential-dependent transmission was blocked with tetrodotoxin (TTX, 1 μM), and ontogenetically-evoked release was rescued by subsequent application of voltage-gated channel blocker 4-aminopyridine (4-AP, 100 μM). To confirm the glutamatergic nature of EPSCs, DNQX (10 μM) was bath applied.

Whole cell patch clamp recordings were obtained using a Multiclamp 700B amplifier (Molecular Devices, California, USA), low-pass filtered at 1 kHz and digitized at a sampling rate of 20 kHz using Digidata 1440 A (Molecular Devices). Data was recorded on a PC using pClamp 10.6 (Molecular Devices) and analyzed using Clampfit (Molecular Devices).

#### Post-recording histology

After recordings, slices were fixed in 4% PFA, washed in PBS within 24 h, and stored in an ethylene glycol–based cryoprotectant at -20°C until processing (<2 weeks). Biocytin-filled neurons were labeled with streptavidin–Alexa Fluor 647 (S21374, Invitrogen, Thermo Fisher Scientific) and imaged on a confocal microscope (STELLARIS 5, Leica Microsystems) using 10× (PL FLUOTAR 10×/0.3) and 63× (APO 63×/1.4 oil immersion) objectives.

### Modelling Methods

We constructed networks of leaky-integrate-and-fire (LIF) models to analyze the potential sources of the up-state and home cage dynamics for calcium activity in CRH neurons ensembles. These networks took the general form of:

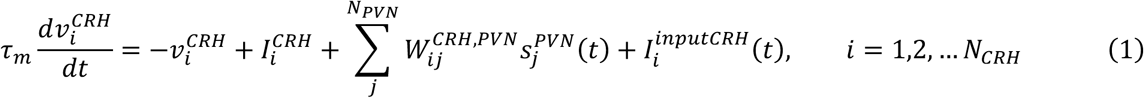

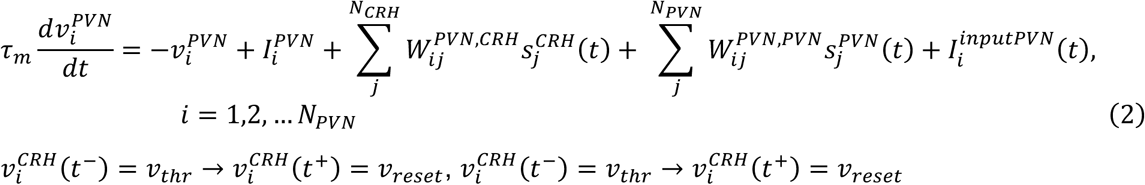

with identical neuronal parameters for all neurons (see Table 1 For a description of parameters). The network consists of two coupled populations, representing the CRH neurons in the PVN (Equation (1)) and a GABAergic population in the peri-PVN/Lateral Hypothalamus area. These populations were coupled by the synaptic weights ***W***^***CRH,PVN***^, ***W***^***PVN, CRH***^, and ***W***^***PVN, PVN***^. These synaptic weights were either randomly drawn as in the case of the disinhibition model (see Supplementary Methods) or were all set equal to 0, as in the case of the tracking/slave models. Every time the voltage trace for a neuron 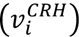 or 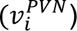 reached the threshold value *v*_*thr*_, this signified the occurrence of a spike, followed by an immediate reset to the voltage *v*_*reset*_. The spikes were filtered with a decay time constant of τ_*d*_ = 20 *ms* and rise time of τ_*r*_ = 2 ms:

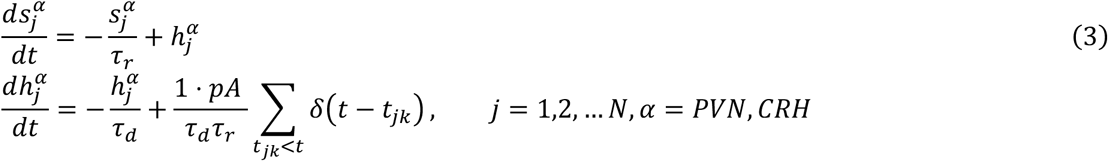

**Table 1.**
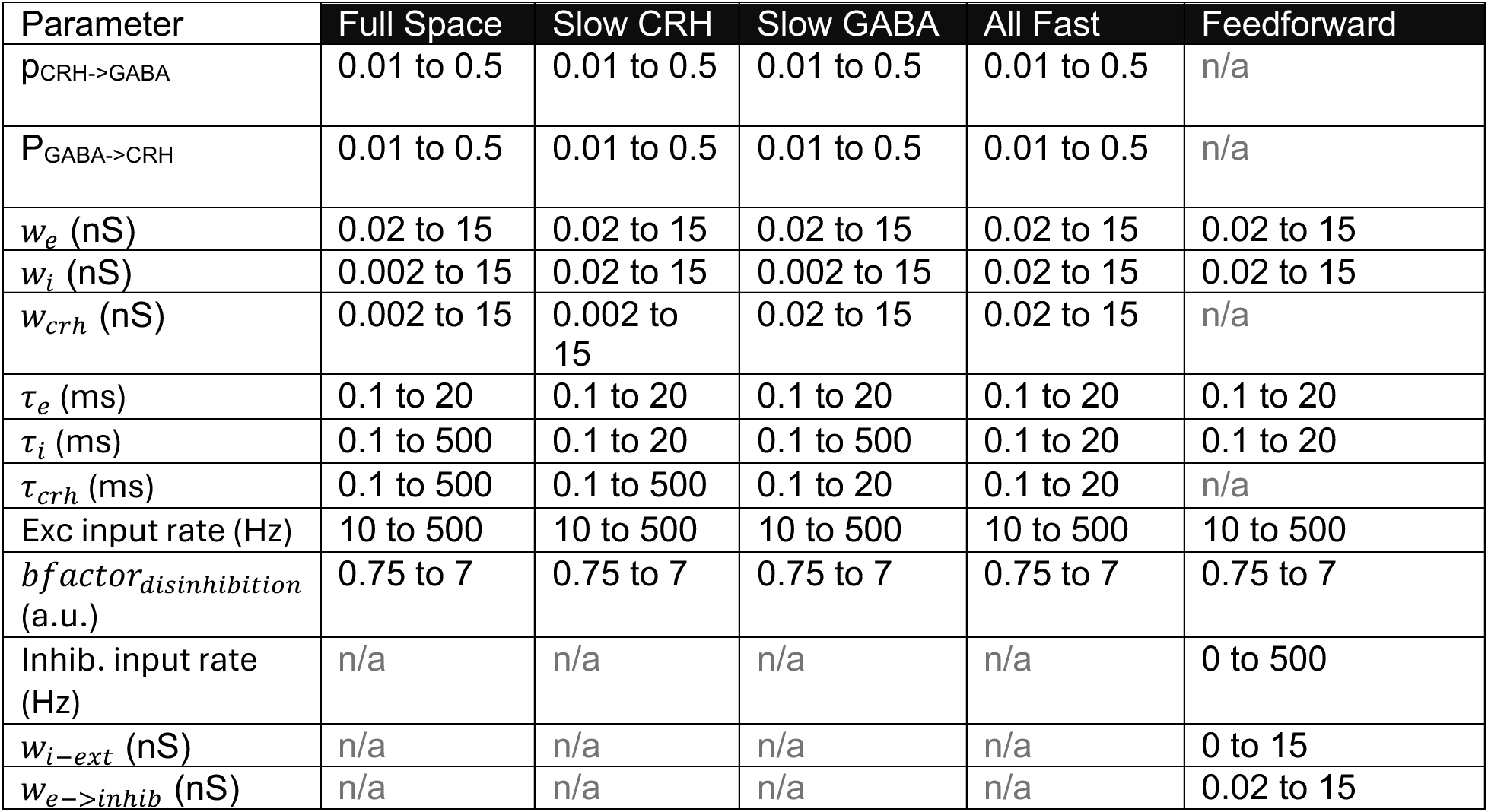
Scan Space for Each Network Motif;

| Parameter | Full Space | Slow CRH | Slow GABA | All Fast | Feedforward |
| --- | --- | --- | --- | --- | --- |
| $p_{CRH \rightarrow GABA}$ | 0.01 to 0.5 | 0.01 to 0.5 | 0.01 to 0.5 | 0.01 to 0.5 | n/a |
| $P_{GABA \rightarrow CRH}$ | 0.01 to 0.5 | 0.01 to 0.5 | 0.01 to 0.5 | 0.01 to 0.5 | n/a |
| $w_e$ (nS) | 0.02 to 15 | 0.02 to 15 | 0.02 to 15 | 0.02 to 15 | 0.02 to 15 |
| $w_i$ (nS) | 0.002 to 15 | 0.02 to 15 | 0.002 to 15 | 0.02 to 15 | 0.02 to 15 |
| $w_{crh}$ (nS) | 0.002 to 15 | 0.002 to 15 | 0.02 to 15 | 0.02 to 15 | n/a |
| $\tau_e$ (ms) | 0.1 to 20 | 0.1 to 20 | 0.1 to 20 | 0.1 to 20 | 0.1 to 20 |
| $\tau_i$ (ms) | 0.1 to 500 | 0.1 to 20 | 0.1 to 500 | 0.1 to 20 | 0.1 to 20 |
| $\tau_{crh}$ (ms) | 0.1 to 500 | 0.1 to 500 | 0.1 to 20 | 0.1 to 20 | n/a |
| Exc input rate (Hz) | 10 to 500 | 10 to 500 | 10 to 500 | 10 to 500 | 10 to 500 |
| $bfactor_{disinhibition}$ (a.u.) | 0.75 to 7 | 0.75 to 7 | 0.75 to 7 | 0.75 to 7 | 0.75 to 7 |
| Inhib. input rate (Hz) | n/a | n/a | n/a | n/a | 0 to 500 |
| $w_{i-ext}$ (nS) | n/a | n/a | n/a | n/a | 0 to 15 |
| $w_{e \rightarrow inhib}$ (nS) | n/a | n/a | n/a | n/a | 0.02 to 15 |

The total synaptic current is given by multiplying the synaptic filtered variables ***S***^***PVN***^(***t***) ***and S***^***CRH***^(***t***) by the appropriate, dimensionless weight matrices in Equation (1) and (2).

The input current 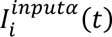, for α = *PVN*, *CRH* was selected to elicit the sparse bursting in the home-cage and a plateau in the up-state and varied from model to model (see later).

To elicit a calcium response, we filtered the CRH spikes with an identical filtering equation as in Equation (3), however we incorporated longer decay and rise times of τ_*D*[*Ca*]_ = 800 ms and τ_*R*[*Ca*]_ = 20 ms based on the parameters from Chen *et al*^42^. All equations were integrated with a forward Euler method with a time-step of 0.05 ms. All simulations were run for a total of 900 seconds for the purposes of comparison with the experimental data. In all networks, we randomly selected a subset of 40 CRH neurons to approximately match the sample sizes obtained via the experimental data.

### Additional Data Analysis Methods

#### Singular Value Decomposition

To determine how the dimensionality of the calcium traces changed due to the exposure of the novel environment, we employed the Singular Value Decomposition to analyze the miniscope data. The singular value data decomposes the traces from the miniscope data, *r*_*i*_(*t*), *i* = 1,2, … *N* as a linear combination in an orthogonal basis set, *u*_j_(*t*), *j* = 1,2, & *N* such that

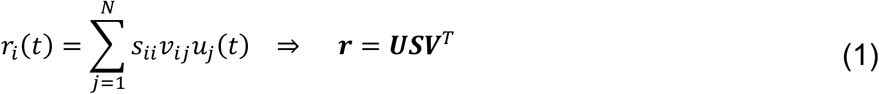

Where *s*_*ii*_ are the *N* singular values, and *v*_*i*j_ is the coefficient of the *i*th trace on the *j*th basis. By using k basis functions, we can find the optimal approximation of our data with a low-rank, or equivalently low-dimensional basis set:

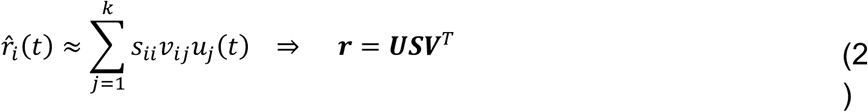

To ascertain the dimensionality of the neural data, we varied *k* and computed the low rank approximation ***r̂*** for each *k* = 1,2, & *N*, and then subsequently computed the percentage error in the approximation as:

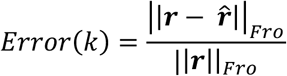

where ||***x***||_*Fro*_ is the Frobenius norm of the matrix x. The dimensionality was estimated as the smallest value of *k* that resulted in a less than 10% reconstruction error. Note that the cutoff value could have been taken at any % reconstruction accuracy. However, as the error curve for the up-state lies to the right of the error curves in the home cage (except at *k* = 0 and *k* = *N*), the up-state always requires more basis elements to reconstruct with an identical level of accuracy as the home-cage.

In general, we found that the first two basis elements *u*_1_(*t*) and *u*_2_(*t*) corresponded to the background activity (tonic), and the dominant fluctuations (phasic) off of the background activity. The second basis vector was tightly correlated to the synchronized bursts in the home-cage both before and after introduction into a novel environment. We sorted the calcium traces according to the magnitude of the basis coefficient *v*_*i*2_, *i* = 1,2, … *N*. The ordered pairs of coefficients (*v*_*i*1_, *v*_*i*2_) did not display any clustering around discrete modes, and thus the neural responses indicate a continuum of behaviours rather than discrete subpopulations, hence the choice of sorting rather than clustering responses.

#### Normalizing Calcium Traces

To better visualize the synchronized responses during the home-cage bursts, we normalized the activity traces prior to plotting the heat-map responses with an *L*_2_ normalization:

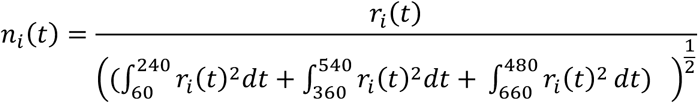

The normalized *n*_*i*_(*t*) better visualize the synchronized responses during home-cage bursts by accounting for neuron-to-neuron differences in the basic calcium level.

#### Synchrony Analysis

To assess the probability of burst coherence occurring randomly simply due to the neuronal firing rates, we applied a series of random time shifts to each calcium trace in each animal. Each neuron was randomly shifted 10^5^ times with a uniform random time shift on the interval [−20,20] s. For each trial, we recomputed the mean calcium signal allowing us to create a null distribution for burst coherence at the population level. A simple Bonferroni adjustment was made to set the critical p-value at 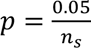 where *n* = 1799 is the number of sample points in each time interval.

#### ROI Analysis

To determine if any nearest-neighbour effects occurred between cells, we computed the Pearson correlation coefficient between the nearest neighbours as measured by the Euclidean distance in the Region of Interests (ROIs) which define a Calcium trace. We then shuffled the ROI indices and subsequently used a kernel density estimator with a bandwidth parameter of 0.05 on the nearest-neighbour correlation coefficients for both the unshuffled and normal densities for the purposed of plotting. If a pair of ROIs were each their nearest neighbour, only one data point was used in computing the resulting density function. To assess statistical significance, we used a two-sample Kolmogorov-Smirnov test on the raw data points, as implemented by the kstest2 function in MATLAB 2019.

## Supplementary Methods for Each Model

### Tracking Model 1

In this model, the network consists of *N*^*CRH*^ = 400 neurons, with no GABAergic PVN neurons. Thus, the network contains no recurrent connections as all non-input weights are 0. The inputs to the CRH neurons consist of the following:

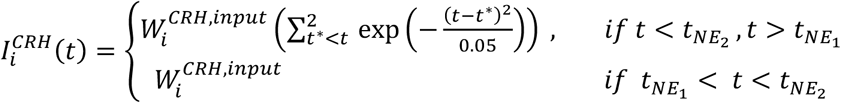

Where 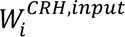 was a randomly drawn synaptic weight from a normal distribution with mean 0 and standard deviation 1. The background current to all neurons was 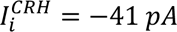. The timepoints *t*_*NE*1_ = 300 and *t*_*NE*2_ = 600 signify the transition to the novel environment, and return to the home cage, respectively. The burst times *t*^∗^ were generated with a uniform distribution over the time intervals [0, *t*_*NE*1_] and [*t*_*NE*2_, *T*] where *T* = 900 *s*, with 10 bursts per interval.

### Tracking Model 2

In this model, we considered the hypothesis that two separate sources of inputs with differing weight strengths elicited the home cage bursts and novel environment up-state, respectively.

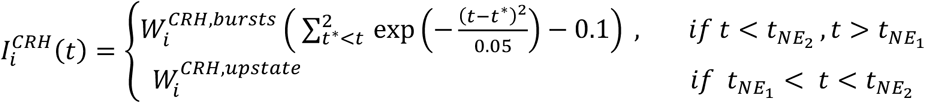

The synaptic weights for the bursts input, 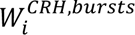 were drawn as in Tracking Model 1. The synaptic weights for the upstate input 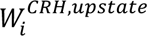 were drawn from a uniform random variable on the interval [0,1]. All other parameters are identical as in Tracking Model 1.

### CRH PVN Network Model with Direct Excitation of CRH Neurons

Unlike the tracking models, we now consider separate populations of PVN and CRH neurons (*N*^*PVN*^ = 200, *N*^*CRH*^ = 200). We assume that the CRH neurons were exclusively excitatory while the other PVN neurons were inhibitory. The weight matrices ***W***^***CRH,PVN***^ and ***W***^***PVN, PVN***^ were randomly generated with 10% sparse connectivity, and each non-zero connection was drawn from a uniform distribution on [-0.1,0]. The excitatory connections ***W***^***PVN, CRH***^were also randomly generated with 1% sparse connectivity with non-zero connections drawn from a uniform distribution on [0,0.5]. The input to the CRH neurons was

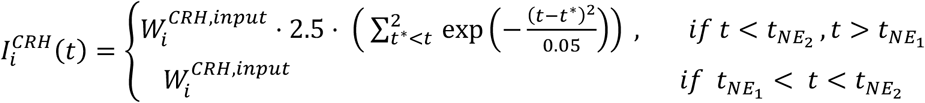

with 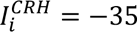, for all *i* = 1,2, … *N*^*CRH*^. The background currents to the CRH neurons was 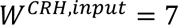 pA while the background current to the PVN neurons was 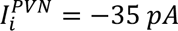

### CRH PVN Network Model with Disinhibition of CRH Neurons

All parameters were identical to the previous implementation, only now the inputs were inhibitory to the GABAergic PVN neurons:

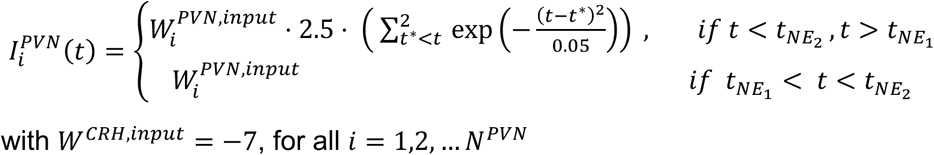

## Spiking Network Model for electrophysiology data

Each network model consisted of a population of 500 CRH and 500 GABA neurons connected recurrently, with varying parameters depending on the motif. Individual neurons were simulated as point neuron models using a modified conductance-based adaptive exponential integrate- and-fire model. ^43^. In short, the membrane voltage, v, Evolves following the equations:

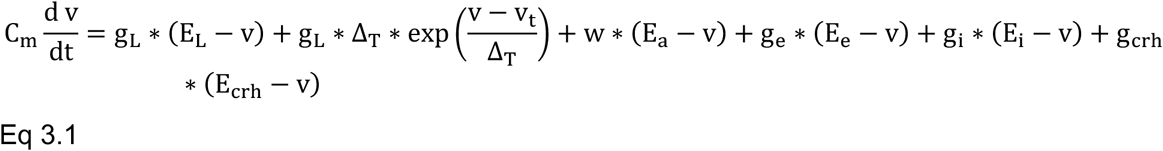

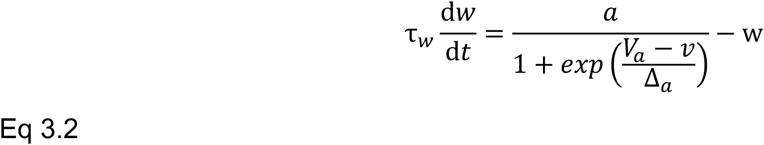

Where, in equation 3.1, *C*_*m*_ is the cell membrane capacitance in pF, *g*_*L*_ is the leak conductance in nS, *E*_*L*_ is the leak equilibrium potential in mV, Δ_Y_ is the exponential slope term in mV, v_Z_is the spike threshold in mV, w is the adaptation conductance in nS, E_[_is the adaptation equilibrium in mV. For the synaptic parameters, *g*_*e*_ is the excitatory conductance in nS, *E*_*e*_ is the reversal potential of the excitatory conductance, *g*_*i*_ is the inhibitory conductance in nS, *E*_*i*_is the reversal potential of the excitatory conductance, *g*_c*r*ℎ_ represents the conductance at the CRH_PVN_->GABA synapse in nS, and *E*_c*r*ℎ_ represents the reversal potential of the CRH conductance.

In equation 3.2, w is the adaptation conductance in nS, τ_a_ is the adaptation time constant in ms, *a* is the maximal conductance in nS, *V*_*a*_ is the subthreshold activation potential mV, and Δ_*a*_is the slope of activation in mV. Following a spike, where *v* > *v*_c*ut*_ we increment according to:

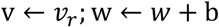

Single neuron model parameters were fit to *ex vivo* patch-clamp electrophysiology recordings as previously described in ^12^. Here, we built a library of parameters based on several fittings, which were up-sampled to populate our network.

The synaptic conductance noted above, evolved according to the following equations:

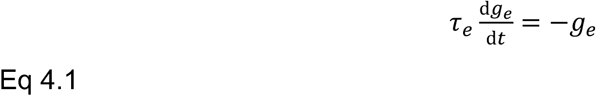

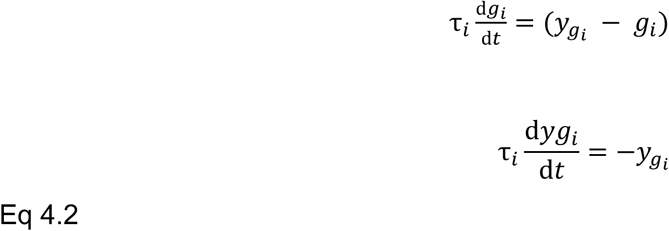

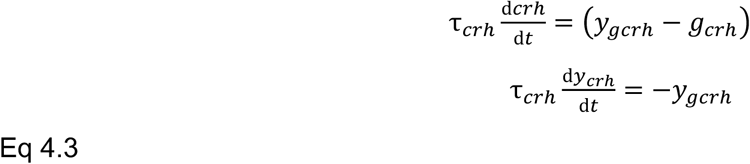

Where τ_*e*_ is the excitatory decay time constant in ms, τ_*i*_is the inhibitory decay time constant in ms, τ_c*r*ℎ_ is the *g*_c*r*ℎ_ decay time constant in ms, −*y*_e*i*_ and −*y*_ec*r*ℎ_ represents the growth offset variables (in nS) for the *g*_*i*_ and *g*_c*r*ℎ_respectively. Upon presynaptic spike, the variables are updated as follows:

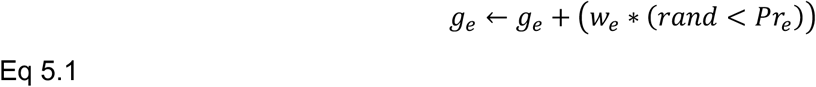

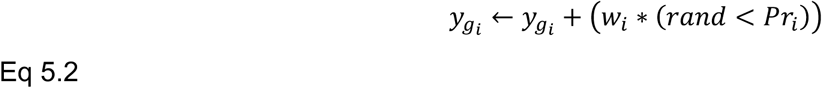

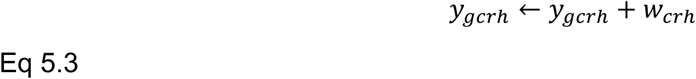

Where *w*_*e*_ is the excitatory synaptic weight in nS, *Pr*_*e*_ is the excitatory synaptic probability of release, *w*_*i*_ is the inhibitory synaptic weight in nS, *Pr*_*i*_is the inhibitory synaptic probability of release, and *w*_c*r*ℎ_ is the CRH-GABA excitatory synaptic weight in nS. In cases where a ‘fast’ synapse was desired for *g*_*i*_ and *g*_c*r*ℎ_ the synaptic variables were incremented directly.

Across all network motifs, disinhibition were driven by a drop in inhibitory probability of release (*Pr*_*i*_), and an increase in spike-triggered adaptation (b). Naturalistic return from disinhibition was modelled using the following equations:

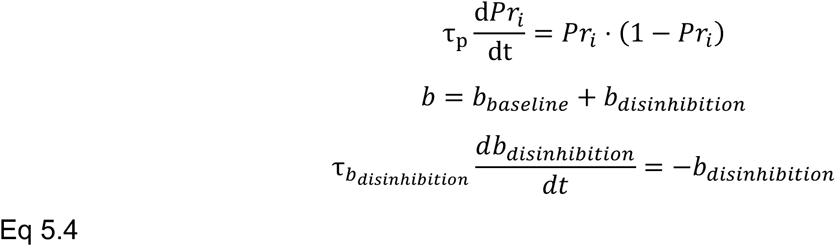

Where *b*_*p*_ is the time constant of the *Pr*_*i*_ decay back to 1, *b*_*disinhibitition*_ is the strength of the increased spike-triggered adaptation in nS, and *b*_*disinhibitition*_ is the time constant of the *b*_*disinhibitition*_ decay in ms. *b*_*disinhibitition*_ was computed for each neuron as *b*_*disinhibitition*_ = ^*b*^_*baseline*_ ∗ *b_disinhibitition_*

To understand the viable parameter space & network motifs, first, a randomized 3600-point scan was run across the entire parameter range. Next, for each network motif presented, a more focused scan was run across a refined parameter space (see Table 1). Each scan consisted of 2400 different search points. To quantify the rudimentary goodness-of-fit of each potential model, each model’s baseline was compared to *in vivo* data as outlined in ^12^. First, a representative spike train was chosen from the *in vivo* data. Then, for each CRH neuron in the network model, the 2d Wasserstein distance between the joint ISI distribution of the neuron and the representative *in vivo* spike train was computed. Next, the lowest 10% of distances from the potential model were averaged and used as the model’s ‘error score’ to implement heterogeneity in the network. Then, for each network motif, the ten lowest error configurations were compared against a second set of baseline *in vivo* features: inter-burst-interval (IBI), probability to burst based on preceding isi, and burst event length. For each network motif, the parameter set that best captured these features was selected as the putative model (see Table 2).

**Table 2.** Optimal Parameter Set for Each network motif;

| Parameter | Slow CRH | Slow GABA | All Fast | Feedforward |
| --- | --- | --- | --- | --- |
| $p_{CRH \rightarrow GABA}$ | 0.3 | 0.1 | 0.01 | n/a |
| $P_{GABA \rightarrow CRH}$ | 0.2 | 0.02 | 0.1 | n/a |
| $w_e$ (nS) | 12.124 | 5.635 | 10.533 | 10.321 |
| $w_i$ (nS) | 2.370 | 11.473 | 13.148 | 5.164 |
| $w_{crh}$ (nS) | 0.0242 | 0.452 | 1.472 | n/a |
| $\tau_e$ (ms) | 14.959 | 12.0928 | 11.874 | 5.017 |
| $\tau_i$ (ms) | 9.404 | 27.0912 | 7.99 | 18.043 |
| $\tau_{crh}$ (ms) | 75.198 | 9.904 | 9.554 | n/a |
| Exc input rate (Hz) | 23.929 | 14.5084 | 15.91 | 241.801 |
| $bfactor_{disinhibiton}$<br>(a.u.) | 6.259 | 3.743 | 4.493 | 2.738 |
| Inhib. input rate (Hz) | n/a | n/a | n/a | 167.592 |
| $w_{i-ext}$ (nS) | n/a | n/a | n/a | 7.829 |
| $w_{e \rightarrow inhib}$ (nS) | n/a | n/a | n/a | 10.321 |

Model performance was evaluated as follows. To examine baseline firing activity (firing rate and CV2), each model was initialized in a low-activity state, followed by disinhibition at 30 seconds, after which activity returned to the starting state with *b*_g_(15 sec) and *b*_*disinhibitition*_ (20 sec, see Eq. 4.1, 4-2): this design mimiced naturalistic fluctuations of activity states observed in vivo under baseline conditions. To mimic the prolonged disinhibition induced by chemogenetic inhibition of ^PVN→^GABAergic neurons, a separate simulation was run in which disinhibition was induced at 5 seconds. In this case, *b*_g_ and *b*_*disinhibitition*_ were set as infinite to capture the sustained effect of chemogenetic inhibition. For both types of simulations, the initial period (5 sec) was excluded from analysis to allow the network to reach a steady state. Firing activities was then averaged across all 500 CRH neurons.

## Competing interests

Authors have no competing interests to declare.

## Notes

### Competing Interest Statement

The authors have declared no competing interest.

### Funding

- CIHR: PJG178289

